# Vertical inheritance governs biosynthetic gene cluster evolution and chemical diversification

**DOI:** 10.1101/2020.12.19.423547

**Authors:** Alexander B. Chase, Douglas Sweeney, Mitchell N. Muskat, Dulce Guillén-Matus, Paul R. Jensen

**Affiliations:** Center for Marine Biotechnology and Biomedicine, Scripps Institution of Oceanography, University of California, San Diego, California; Marine Biology Research Division, Scripps Institution of Oceanography, University of California, San Diego, California

**Author notes:** Corresponding author: AB Chase.

**Keywords:** *Salinispora*, homologous recombination, microbial ecology, speciation

## Abstract

While specialized metabolites are thought to mediate ecological interactions, the evolutionary processes driving their diversification, particularly among closely related lineages, remain poorly understood. Here, we examine the evolutionary dynamics governing the distribution of natural product biosynthetic gene clusters (BGCs) using 118 strains within the marine actinomycete genus *Salinispora*. While previous evidence indicated that horizontal gene transfer (HGT) largely contributed to BGC diversity, we find that a majority of BGCs in *Salinispora* genomes are conserved through processes of vertical descent. In particular, vertical inheritance maintained BGCs over evolutionary timescales (millions of years) allowing for BGC diversification among *Salinispora* species. By coupling the genomic analyses with high-resolution tandem mass spectrometry (LC-MS/MS), we identified that BGC evolution in *Salinispora* proceeds largely through gene gain/loss events and constrained recombination that contributes to interspecies diversity at the gene, pathway, and metabolite levels. Consequently, the evolutionary processes driving BGC diversification had direct consequences for compound production and contributed to chemical diversification, as exemplified in our case study of the medically relevant proteosome inhibitors, the salinosporamides. Together, our results support the concept that specialized metabolites, and their cognate BGCs, represent functional traits associated with niche differentiation among *Salinispora* species.

**GRAPHICAL ABSTRACT:** 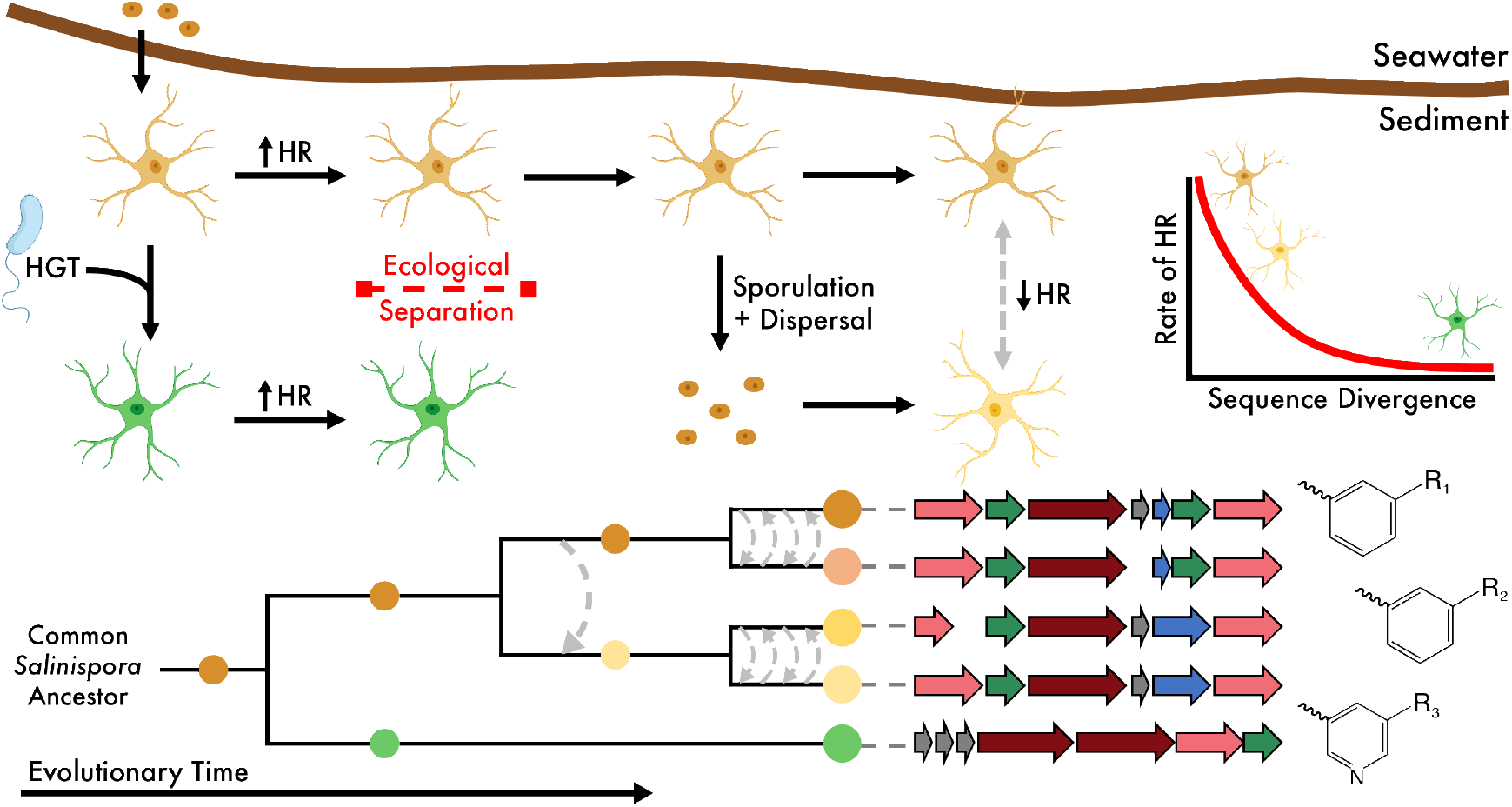

SIGNIFICANCE
Natural products are traditionally exploited for their pharmaceutical potential; yet what is often overlooked is that the evolution of the biosynthetic gene clusters (BGCs) encoding these small molecules likely affects the diversification of the produced compounds and implicitly has an impact on the compounds’ activities and ecological functions. And while the prevailing dogma in natural product research attributes frequent and widespread horizontal gene transfer (HGT) as an integral driver of BGC evolution, we find that the majority of BGC diversity derives from processes of vertical descent, with HGT events being rare. This understanding can facilitate informed biosynthetic predictions to identify novel natural products, in addition to uncovering how these specialized metabolites contribute to the environmental distribution of microbes.

---

While specialized metabolites are thought to mediate ecological interactions, the evolutionary processes driving their diversification, particularly among closely related lineages, remain poorly understood. Here, we examine the evolutionary dynamics governing the distribution of natural product biosynthetic gene clusters (BGCs) using 118 strains within the marine actinomycete genus *Salinispora*. While previous evidence indicated that horizontal gene transfer (HGT) largely contributed to BGC diversity, we find that a majority of BGCs in *Salinispora* genomes are conserved through processes of vertical descent. In particular, vertical inheritance maintained BGCs over evolutionary timescales (millions of years) allowing for BGC diversification among *Salinispora* species. By coupling the genomic analyses with high-resolution tandem mass spectrometry (LCMS/MS), we identified that BGC evolution in *Salinispora* proceeds largely through gene gain/loss events and constrained recombination that contributes to interspecies diversity at the gene, pathway, and metabolite levels. Consequently, the evolutionary processes driving BGC diversification had direct consequences for compound production and contributed to chemical diversification, as exemplified in our case study of the medically relevant proteosome inhibitors, the salinosporamides. Together, our results support the concept that specialized metabolites, and their cognate BGCs, represent functional traits associated with niche differentiation among *Salinispora* species.

Despite linkages between abiotic factors and bacterial diversity (1–3), the key functional traits driving biotic interactions in microbial communities remain poorly understood. These functional traits likely include specialized metabolites, or small molecule natural products, that are known to modulate ecological interactions between organisms (4). Specialized metabolites, which include molecules to known as act as antibiotics and siderophores, can drive biotic interactions via mechanisms such as competition, nutrient uptake, and defense. Taken together, the functions of these small molecules likely represent a major driver of microbial community composition (4). Given that microbes produce a wide variety of biologically active natural products, these compounds may contribute to the diversification and environmental distribution of microbes, as has been observed in eukaryotes (5). To date, however, studies of bacterial specialized metabolite production have largely focused on the discovery of compounds with pharmaceutical potential as opposed to understanding their ecological and evolutionary significance.

Marine sediments represent a unique environment to assess the relationships between microbial diversity and specialized metabolite production. Sediments are associated with diverse microbial communities (6) where competition for limited resources is facilitated via the secretion of small molecules (7, 8). Among bacteria inhabiting marine sediments, actinomycetes such as the genus *Salinispora* are well-known for the production of specialized metabolites (9-11). *Salinispora* (family: Micromonosporaceae) is a member of the rare biosphere in surface sediments (12) and can readily be cultured from tropical and sub-tropical locations (13). This relatively recently diverged lineage includes nine closely related species that share >99% 16S rRNA sequence similarity (14). The presence of this “microdiversity” suggests that fine-scale trait differences contribute to differential resource utilization and niche partitioning (15, 16). Indeed, two *Salinispora* species were recently shown to employ differential ecological strategies for resource acquisition (17), providing insights into the ecological and evolutionary mechanisms contributing to *Salinispora* diversification.

While *Salinispora* has proven a robust model for natural product discovery (18), much remains to be resolved concerning the evolutionary dynamics of the biosynthetic gene clusters (BGCs) encoding these compounds. Prior analyses of *Salinispora* BGCs revealed extensive horizontal gene transfer (HGT) and exchange both within and between species (19), suggesting a “plug and play” model of BGC evolution (20). These observations are consistent with the prevailing view in natural product research that BGCs are rapidly gained and lost via HGT (21-23), particularly within Actinobacteria taxa (24). In *Salinispora*, the exchange of BGCs between species is likely mediated by conjugative elements, as in Streptomyces (25), and further facilitated by the absence of geographic barriers in their distribution (26). At the same time, profiles for the production of specialized metabolites encoded by *Salinispora* BGCs revealed species-specific patterns (27), providing competing models for BGC evolution.

Resolving the evolutionary dynamics driving BGC distributions in *Salinispora* can help inform future natural product discovery efforts. By one account, the horizontal exchange of *Salinispora* BGCs may be frequent and highly dependent on the local community (19, 20), resulting in similar strains from different locations yielding different metabolites. Alternatively, the small molecules encoded by BGCs may represent phylogenetically conserved traits (27, 28), in which case similar strains produce similar metabolites. To reconcile these seemingly contrasting scenarios, we sought to identify the evolutionary processes contributing to the distribution of BGCs. We hypothesized that specialized metabolite production is subject to strong selective pressures, with compounds and their associated gene clusters representing functional traits contributing to niche differentiation. If correct, we expected the distribution of BGCs, regardless if they were horizontally acquired at some point in time (19), to be subsequently maintained within *Salinispora* through processes of vertical descent. To better understand the role of vertical inheritance in driving BGC evolution, we revisited the distribution of BGCs in 118 genomes across the nine newly described *Salinispora* species (14). Finally, to assess the functional consequences of BGC diversification on compound production, we focused on nine experimentally characterized BGCs and applied targeted tandem mass spectrometry to detect their associated molecules. Our results support the hypothesis that specialized metabolites represent functional traits contributing to ecological differentiation among closely related *Salinispora* species.

## RESULTS

### *Salinispora* delineated by biosynthetic potential

Our molecular clock analysis indicated that *Salinispora* recently diverged within the Micromonosporaceae family 89.1±37.1 million years ago (MYA; Figure S1); yet the genus has already differentiated into nine species (Figure 1A). To gain insights into the functional traits that promoted differentiation within *Salinispora*, we first investigated differences in gene content among the 118 *Salinispora* genomes (a.k.a., the flexible genome). Flexible gene composition was highly congruent with species designations (Figure 1B), with strains within the same species sharing more flexible genes than expected by chance and explained 53.1% of the variation in the flexible genome (permutational multivariate analysis of variance (PERMANOVA); p<0.01). Geographic location, to a lesser degree, accounted for 8.3% of the variation in flexible gene composition (p<0.01). Within the flexible genome, we also identified species-specific orthologs, genes shared by all strains within a species but not observed in any other *Salinispora* species. These genes, which should encode the functional traits that define *Salinispora* species, were largely annotated as hypothetical proteins (Figure S2A). Of the available annotations, 18.1±15.3% of the species-specific orthologs were associated with specialized metabolism (Figure S2B). In addition, when we searched the genomic regions flanking all species-specific orthologs, regardless of annotation, we found that 28.9±26.2% were located within the boundaries of predicted biosynthetic gene clusters (BGCs; Figure S2C).

**FIGURE 1.**
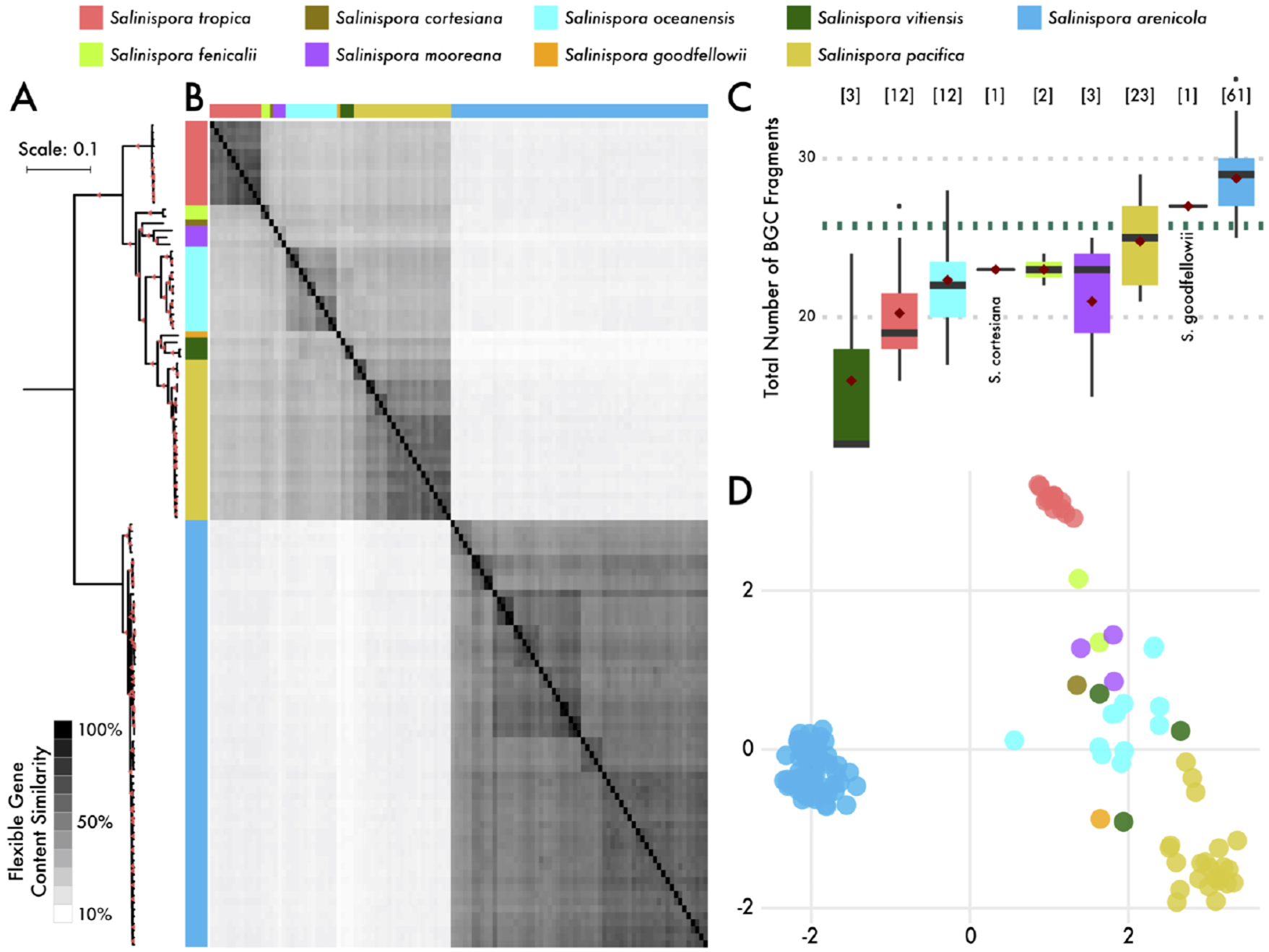
Genetic relatedness of globally distributed *Salinispora* strains (N=118). **A)** Core genome phylogeny based on core genes. Colors denote species. Bar, 0.1 nucleotide substitutions per position. Bootstrap values indicate >90% support. **B)** Flexible gene content similarity between strains. Heatmap was generated from a Jaccard distance matrix. **C)** Total number of biosynthetic gene cluster (BGCs; both whole and fragments) identified across *Salinispora* species. Black bar represents median distribution, red diamond represents mean distribution. Dashed line denotes genus average. [N] represents number of genomes per species. **D)** Non-metric multidimensional scaling (NMDS) plot depicting gene cluster family (GCF) composition for each *Salinispora* strain. Circles represent strains colored by species.

Given the high percentage of species-specific flexible genes associated with specialized metabolism, we expected that BGC diversity and distribution would similarly correspond with *Salinispora* species diversity. To address this, we identified a total of 3041 complete or fragmented (on contig edges) BGCs across all 118 *Salinispora* genomes (mean = 25.8 per genome) accounting for 18±2.3% of an average 5.6 Mbp *Salinispora* genome. When compared to other bacterial genera, including taxa well-known for specialized metabolite production (e.g., *Moorea* and *Streptomyces), Salinispora* dedicated the highest genomic percentage to this form of metabolism (Figure S1), further highlighting its importance in this genus. Between *Salinispora* species there was significant variation in the total number of BGCs (Figure 1C; analysis of variance [ANOVA]; p<0.001) and the genomic percentage dedicated to specialized metabolite production (Figure S1; ANOVA; p<0.001). In cases where the number of genome sequences are low (e.g., *S. vitiensis)*, BGC abundances may not be representative of the species.

To compare BGC composition across species, the 3041 predicted BGCs were grouped into 305 gene cluster families (GCFs; Figure S3A). Similar to prior reports (20), 35% of the GCFs were populated by a single BGC (Figure S3B). It can be inferred that these BGCs represent relatively recent acquisition events that are not well represented in our genomic dataset. Despite representing a large percentage of the total GCF diversity, the singleton BGCs comprised only 3.6% of the BGCs detected among all strains (108 out of 3041) and equate, on average, to only 0.9 BGCs per *Salinispora* genome (inset Figure S3B). In contrast, the vast majority of BGCs were shared among strains (Figure S3C). As in flexible gene content, we found that 43.6% of the variation in GCF composition was explained by species designation (Figure 1D; PERMANOVA; p<0.01), with geography explaining an additional 11.1% (p<0.01). Correlations between BGC distributions and species delineations are further supported by BGC average nucleotide identity (ANI) values, which were highly similar to the whole-genome ANI values used to delineate species boundaries (Figure S4). Nonetheless, a small percentage of shared BGCs (1.4%) showed evidence of relatively recent interspecific transfers (Figure S4B). Collectively, these results indicate that HGT events provide a mechanism to expand BGC diversity, while at the same time, BGC composition is largely driven by processes of vertical descent.

### Drivers of BGC evolution

Given that BGC distributions were largely explained by shared phylogenetic history, we sought to identify the specific evolutionary processes that may contribute to BGC diversification. To do so, we concentrated on nine experimentally characterized BGCs that range in conservation from species-specific to ubiquitous in the genus (Table S2). Indeed, the nine BGCs span a range of evolutionary time, including BGCs that were present prior to *Salinispora* speciation >100 MYA (i.e., *lym, sta*, and *spt*), recently evolved BGCs (i.e., *slc* 6.3-18.3 MYA), and BGCs that are prone to HGT events (Figure S5A). Despite these differences, phylogenies of all nine BGCs (Figure S5B) revealed that genetic differentiation within BGCs remained a function of time as they were maintained by vertical inheritance (Figure 2; multiple r^**2**^=0.84, p<0.001).

**FIGURE 2.**
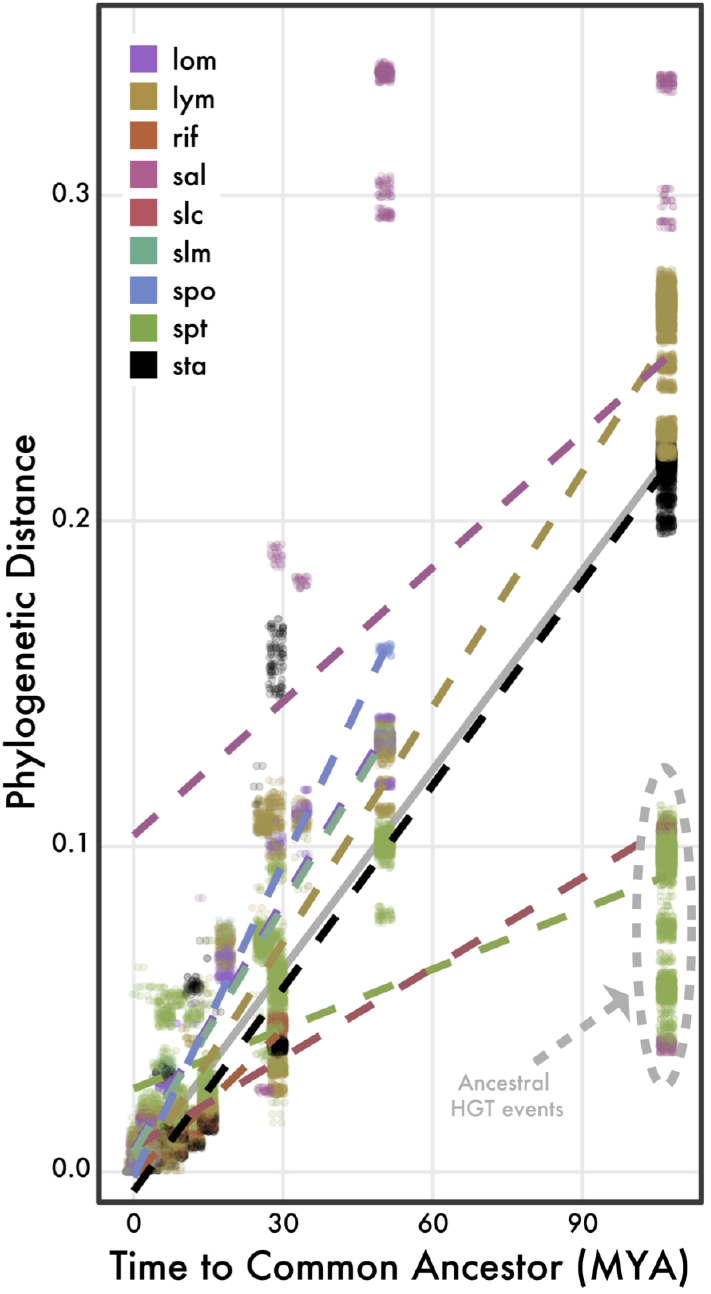
Phylogenetic distance of BGCs as a function of time (millions of years; MYA). Each point represents a pairwise comparison between *Salinispora* strains that share a BGC (colored by BGC) with divergence times for the same strain pairs. Linear regression lines are denoted for each BGC, with expected neutral divergence (based on core genome phylogenetic distance) in gray. BGCs with predicted transfer events between S. *arenicola* and S. *tropica* are circled for *sal, slc*, and *spt*.

The maintenance of BGCs over evolutionary time indicates that other evolutionary processes can contribute to BGC diversification. An eventinference parsimony model indicated that a variety of evolutionary processes, including BGC duplication, transfer, and loss events, contributed to the observed BGC distributions (Table 1). In particular, there were frequent intraspecific recombination events, averaging 18.7±14.7 across the nine BGCs, relative to interspecific horizontal transfers (0.9±1.1), supporting the previous ANI result that horizontal transfer of BGCs between species are rare (Figure S4). In the few cases where interspecies transfers occurred (e.g., *sal* and *slc* BGCs), the BGCs remained monophyletic post-transfer (Figure S5B) and their genetic divergence remained a function of divergence time (Figure 2), supporting a single transfer event followed by vertical inheritance. The model further revealed that BGC loss events (Table 1) explained the patchy distributions of some BGCs. For instance, the *lom* BGC followed a strict model of vertical inheritance with predicted loss events in *S. mooreana* and *S. oceanensis* (Figure S5A). Together, these results indicate that diversification in the nine BGCs is highly correlated to divergence time with frequent intraspecies recombination and loss events.

**Table 1.** Recombination, duplication, transfer, and loss events of nine BGCs relative to the core genome.

| BGC | Num. of Species | Num. of strains | Alignment Length (bp) | Recombination Metrics |  |  |  | DTL Model Predicted Events |  |  |  |
| --- | --- | --- | --- | --- | --- | --- | --- | --- | --- | --- | --- |
|  |  |  |  | R/θ | δ | v | r/m | Interspecific Transfer | Intraspecific Transfer | Duplication | Loss |
| lom | 6 | 42 | 84964 | 0.016 | 6696.0 | 0.026 | 2.77 | 0 | 10 | 0 | 14 |
| lym | 9 | 109 | 32597 | 0.019 | 4101.2 | 0.021 | 1.63 | 0 | 33 | 1 | 33 |
| rif | 1 | 59 | 91550 | 0.655 | 6368.3 | 0.003 | 13.90 | 0 | 23 | 0 | 15 |
| sal | 6 | 30 | 15911 | 0.024 | 3453.9 | 0.032 | 2.70 | 2 | 5 | 0 | 16 |
| slc | 2 | 25 | 31937 | 0.020 | 1316.7 | 0.058 | 1.57 | 1 | 11 | 1 | 5 |
| slm | 3 | 25 | 28111 | 0.189 | 428.0 | 0.041 | 3.28 | 1 | 10 | 0 | 14 |
| spo | 2 | 13 | 167237 | 0.018 | 1032.6 | 0.093 | 1.71 | 1 | 5 | 0 | 3 |
| spt | 7 | 112 | 13368 | 0.904 | 1115.5 | 0.010 | 10.16 | 3 | 49 | 0 | 22 |
| sta | 4 | 77 | 19319 | 0.069 | 1352.4 | 0.032 | 2.96 | 0 | 22 | 1 | 23 |
| Core Genome | 9 | 28 | 4668606 | 0.061 | 1111.6 | 0.026 | 1.75 |  |  |  |  |

R/θ = relative rate of recombination to mutation
δ = mean import length of recombining DNA
v = mean divergence of imported recombining DNA
r/m = relative impact of recombination to mutation accumulation on per-site substitution rate (equal to R/θ \* δ \* v)

To better understand the impact of recombination in structuring the genetic diversity within the nine BGCs, we calculated the ratio at which nucleotides are replaced by either recombination or point mutations *(r/m)*. At the genome level, high levels of recombination (Table 1; *r/m*=1.8) among closely related strains (v=0.03 or 3% genetic divergence among strains) maintains genetic cohesion within species. In fact, a recombination network revealed no recent gene flow between recombining populations within *Salinispora* species (Figure S6) suggesting this genetic isolation may be evidence of nascent speciation events. Similarly, at the BGC level, high levels of recombination (all *r/m*>1.5) were restricted to events between closely related strains (V_MEAN_=3.5%; Table 1). For example, recombination in the *rif* BGC (*r/m*=13.9) was restricted to strains that have only diverged by <0.3%. Recombination events were also restricted in the size of their recombining segments, as the average length of a recombining segment (6) was a fraction of the total BGC length (Table 1). Finally, these events had varying effects on the genetic diversity of the nine BGCs. For instance, the two most widely distributed BGCs, *lym* and *spt*, exhibited drastically different *r/m* values (1.6 and 10.2, respectively). As a result, reduced recombination allowed the *lym* BGC to evolve in accordance with the core genome, while frequent recombination of the *spt* BGC limited its divergence (Figure 2). While recombination can homogenize genetic diversity, these events were limited to small sections of the BGC.

Homologous recombination can also facilitate gene-specific sweeps (29), which can be evident by reduced nucleotide diversity. A comparison of the conserved biosynthetic genes found in the nine BGCs with those found in the core genome revealed reduced nucleotide diversity in the *sal* and *spo* BGCs in *S. pacifica* and the *slc* BGC in *S. arenicola* (Figure S7A). However, the BGCs in these three instances were only observed in a small number of closely related strains (i.e., three *S. pacifica* and five *S. arenicola* strains sharing >99.6% and >99.4% genome-wide ANI, respectively), which likely accounts for the reduced diversity. In contrast, most conserved biosynthetic genes showed no evidence of recent selective sweeps, with the relatively high nucleotide diversity indicating that recombination was insufficient to prevent BGC diversification. Depressed gene flow observed in the recombination network (Figure S6) provides further support that recombination is unlikely to constrain species-level BGC diversification. Notably, this diversification was neutral based on analyses of selection coefficients (dN/dS). Among the 134 conserved biosynthetic genes analyzed, 97.8% had dN/dS<1, indicating neutral, nondirectional selection (Figure S7B). Thus, it appears that the high nucleotide diversity in BGCs, such as *rif* (Figure S7A), is due to ancestral sweep events followed by neutral divergence and constrained recombination among populations.

### Specialized metabolites as functional traits

Since representative BGCs for each of the nine GCFs have been experimentally characterized (Figure S8) (30-38), we next sought to understand how the observed evolutionary dynamics contributing to BGC diversification affected, if at all, production of the compound. By applying untargeted liquid chromatography, high-resolution tandem mass spectrometry (LC-MS/MS) to 30 representative strains across the nine *Salinispora* species, we first detected a total of 3575 unique molecular features from cultured extracts. The total number of metabolites, or the metabolome, revealed that strains within the same species produced more similar molecular features than strains between species (Figure 3A; PERMANOVA; p<0.01), with 44.9% of the variation explained by species designation. These results, in combination with the GCF composition (Figure 1D), provide clear evidence that vertical descent plays a major role in structuring *Salinispora* specialized metabolism.

**FIGURE 3.**
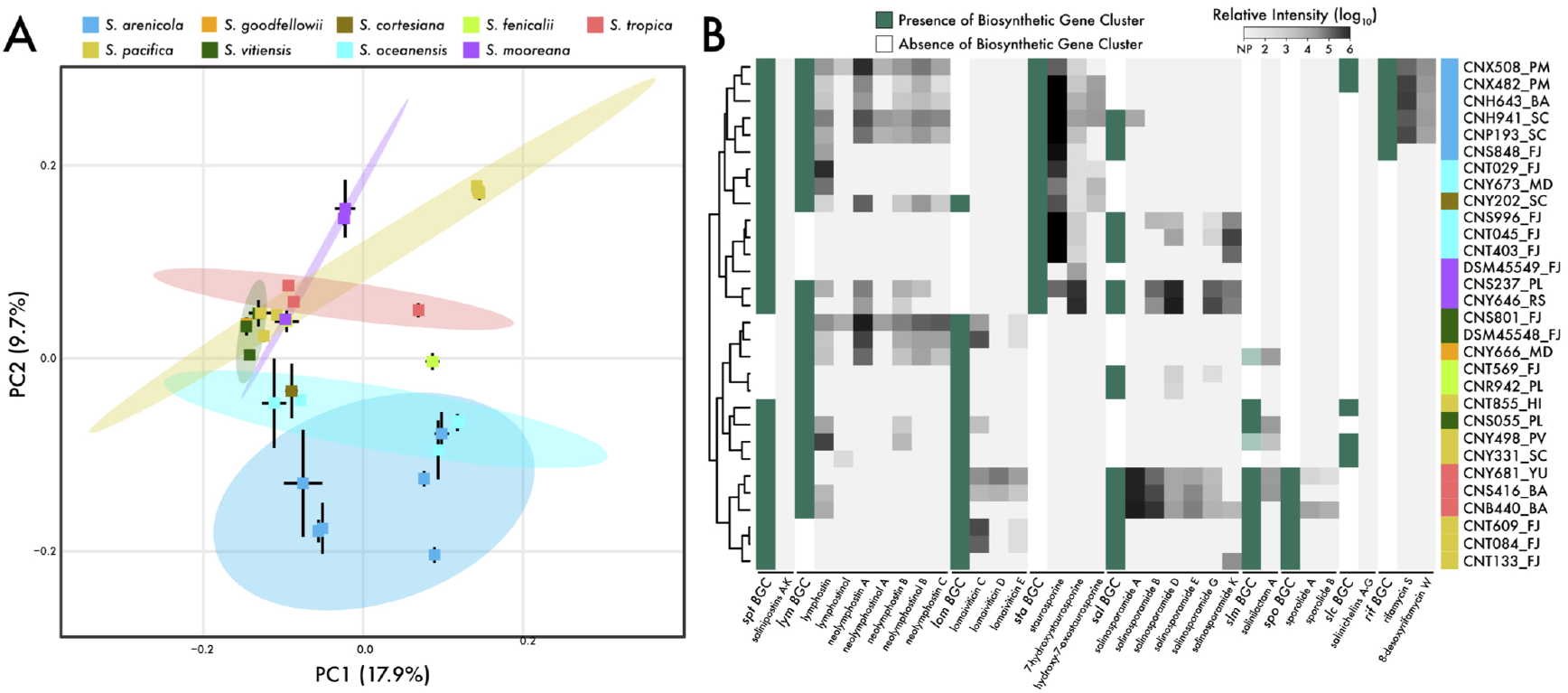
*Salinispora* metabolomics and BGCs distributions across nine species (N=30 strains). **A)** Principal Components Analysis (PCA) showing relationship among strain metabolomes. Each point represents a strain colored by species. Standard deviations shown as black lines for each strain and ellipses as 95% confidence intervals for each species. **B)** Targeted analysis for the products of nine BGCs. Hierarchical clustering of the relative production of analogs generated from a Euclidean distance matrix. Green/white bars show the BGC presence/absence. Light green bars indicate the detection of a BGC fragment. Gray/black bars show the relative production of each identified analog, NP = no production. X-axis lists BGC name followed by product names. Y-axis lists strains colored by species with abbreviations for geographic origin (see Table S1 for more information).

At some level, the genetic diversity observed within GCFs should translate into structural differences in the compounds. Considering the nine BGCs analyzed above, we noted a range of genetic differences from single nucleotide polymorphisms (SNPs) to large variations in gene content. To address how these genetic changes may affect compound production, we applied targeted LC-MS/MS from the culture extracts to detect 25 known compounds and many putative analogs encoded by the BGCs (Table S3). The presence of a BGC did not always equate to product detection (i.e., salinichelins and salinipostins were not detected), suggesting the culture or extraction methods may not have been appropriate. However, the products of seven BGCs were detected, with strains from the same species preferentially producing similar compounds and their known associated analogs (Figure 3B; PERMANOVA, p<0.01). These results indicate that BGC diversification at the species level translated to finer species-specific signatures in terms of analog production.

Importantly, the genetic differences associated with BGC diversification varied in their impact on metabolite production. At the genetic level, the relatively high intraspecies nucleotide diversity observed within the *rif* BGC in *S. arenicola* (Figure S7A) did not reflect differences in rifamycins detected (Figure 3B). This diversity, which was considered neutral in terms of dN/dS values, does not affect compound production as all strains maintain production of the potent antibiotic rifamycin S in *S. arenicola*. Conversely, subtle gene differences in auxiliary enzymes in the sta BGC affected compound production. While three of the four species with this BGC produced detectable staurosporine, *S. mooreana* preferentially produced 7-hydroxystaurosporine (Figure 3B). Comparative genomics revealed that the NAD-dependent dehydratase enzyme is missing in *S. mooreana* strains with the *sta* BGC (Figure S9A), which likely accounts for the presence of the hydroxy group in 7-hydroxystaurosporine produced by this species (Figure S8). More pronounced interspecies polymorphisms were observed in the *spo* BGC between *S. tropica* and the subset of *S. pacifica* (3 of 23 strains) that possess the BGC. While all strains maintain the type I polyketide synthase (PKS) responsible for the polyketide core (38), the three *S. pacifica* strains lack the 45 kbp nonribosomal peptide synthetase (NRPS) region responsible for biosynthesis of the cyclohexenone epoxide subunit (Figure S9B). Correspondingly, the S. pacifica strains did not produce sporolides (Figure 3B) or any other derivatives that could be identified. Interestingly, the other 20 *S. pacifica* strains that lack the *spo* BGC encode a similar enediyne BGC linked to cyanosporaside production (19), suggesting the products may perform similar ecological functions. Together, these results directly link BGC diversification to structural changes in the encoded metabolites.

### Salinosporamides: a case study for BGC evolution

To further illustrate the evolutionary processes driving chemical diversification, we examined the *sal* BGC, which encodes the biosynthesis of the anti-cancer agent salinosporamide A and analogs (30, 39). While salinosporamides A and K were originally reported from *S. tropica* (40) and *S. pacifica* (39, 41), respectively, we now show that the *sal* BGC is observed in six of the nine *Salinispora* species (Figure 4A). Phylogenetic analysis provides evidence that the *sal* BGC was recently transferred 5.4±2.6 MYA between *S. arenicola* and *S. tropica* (Figure 4A) but has otherwise descended vertically within the genus for >50 MYA (Figure S6A). Notably, the *sal* BGC is rapidly diverging between species (note deviation of the *sal* BGC compared to core genome in Figure 2), while at the same time being highly conserved within species (Figure S10). Thus, the *sal* BGC provides a useful model to address the relationships between species diversification, BGC composition, and compound production.

**FIGURE 4.**
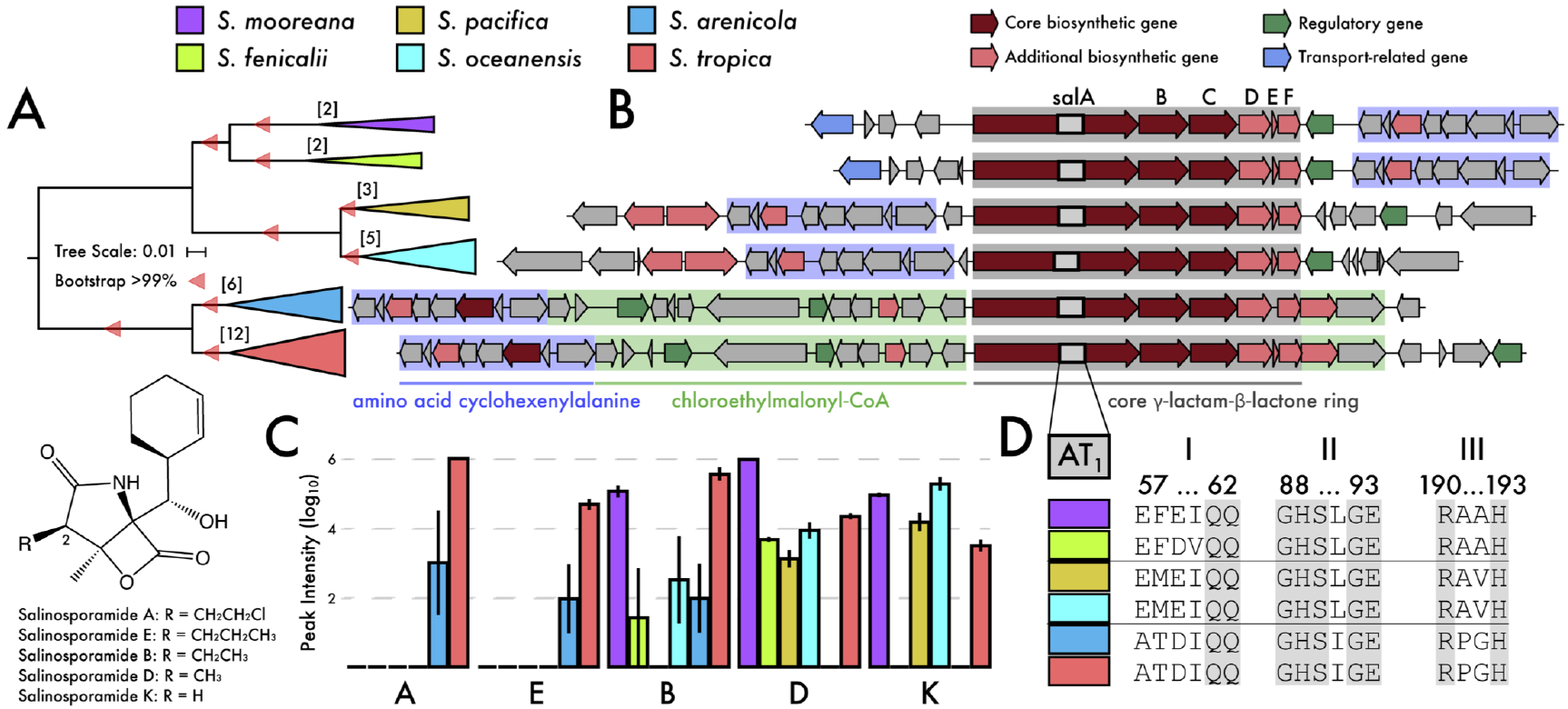
Variations in the salinosporamide BGC *(sal)* and its products. **A)** Phylogeny of the *sal* BGC. Colors denote species. Bar, 0.01 nucleotide substitutions per position. Bootstrap values indicate >99% support. Brackets indicate number of genomes in each species encoding the BGC. **B)** Representative BGC for each species. Genes are colored by predicted biosynthetic function with gene blocks colored by their role in compound production. **C)** Salinosporamides production across species. Analogs are listed in descending order based on R-group side chain size. **D)** Sequence alignments of the *salA* AT_1_ domain showing three signature motifs associated with substrate specificity. Conserved amino acid regions in grey.

We performed targeted metabolomics on 16 strains encoding the *sal* BGC across the six *Salinispora* species (Table S3). All strains maintain the biosynthetic genes responsible for the core y**-**lactam**-**p**-**lactone ring *(salABCDEF)* and the cyclohexenylalanine amino acid residue, although the position of the latter varied (Figure 4B). While 15/16 strains produced detectable amounts of salinosporamides, the analogs and their yields varied across species (Figure 4C; ANOVA, p<0.01). *S. tropica* and *S. arenicola* were the only species in which salinosporamide A was detected and they are the only species to possess the 15kbp region of the BGC responsible for assembly of the chloroethyl-malonyl**-**CoA extender unit (Figure 4B) (30). This is the first report of salinosporamide A production in *S. arenicola*, although the presence of the BGC in this species was previously documented (20, 42). While the *sal* BGCs in *S. tropica* and *S. arenicola* are nearly identical (e.g., *salA* and *salB* genes share 95.1% and 97.5% amino acid sequence identity between species, respectively), the production of salinosporamides in *S. arenicola* was reduced relative to *S. tropica* (Figure 4C; note log scale). This could be due to a splicing event in the overlapping coding regions in *salD/E* (Figure 4B), which may render *salE*, a chaperone MbtH-like protein in NRPS biosynthesis (43), inactive.

Molecular networking provided additional insights into species-specific patterns of salinosporamide production (Figure S11). Here, we detected potentially new salinosporamide derivatives, including one *(m/z* 307.154) produced exclusively by *S. mooreana*. Inversions of the cyclohexenylalanine amino acid encoding region of the *sal* BGC in *S. mooreana* and *S. fenicalii* (Figure 4B) could facilitate novel structural modifications by down-stream auxiliary genes (Figure S10). The network also revealed evidence that salinosporamide analog production varied by species. We investigated this by analyzing the C-2 γ-lactam side chain and found that *S. tropica* and *S. arenicola* produced compounds with the largest R-groups (Figure 4C). This results from the incorporation of chloroethyl-CoA or propyl-malonyl**-**CoA extender units yielding salinosporamides A and E, respectively (30). Similarly, *S. fenicalii* and *S. mooreana* preferentially produce salinosporamides B and D with medium sized R-groups, while *S. oceanensis* and *S. pacifica* preferentially produce salinosporamide K, which has the smallest R-group. Based on these observations, we suspected substrate discrimination by the *salA* PKS acyltransferase domain (AT_1_). Indeed, the three signature motifs related to extended unit specificity (39) were highly conserved at the species level and strongly associated with the length of the C-2 R-group in the salinosporamide product (Figure 4D, Figure S12). Structural protein modelling of the AT_1_ domain identified motif I (amino acid residues 57-59) as a novel extension of the active site that possibly allows for extender unit discrimination (Figure S13A). In this case, the smaller, polar residues (Ala57 and Thr58) in *S. tropica* and *S. arenicola* (Figure S13B) could accommodate larger substrates yielding salinosporamides A and E, while the larger residues observed at these positions in other species may not.

Thus, salinosporamide production and bioactivity (i.e., salinosporamide A is 100x more cytotoxic than salinosporamide K (39)) is likely affected not only by the ability to synthesize the various extender units (30) but also recognition and selection of the appropriate substrate by the AT_1_ domain. In conclusion, understanding the evolutionary processes for the salinosporamides allowed for extensive biosynthetic insights into the genetic polymorphisms that contribute to chemical diversification and differential analog bioactivity.

## DISCUSSION

It has become increasingly clear that the fine-scale genomic diversity observed in microbial communities reflects the large number of ecologically distinct lineages that co-occur within microbiomes (44-46). At the same time, the phenotypic traits that allow for the coexistence of closely related lineages remain less clear (15, 47). By analyzing strains from the recently diverged genus *Salinispora* (>99% similarity in the 16S rRNA gene), we associated interspecies genetic diversity to phenotypic variation in the form of specialized metabolism. The flexible genome, which is responsible for niche and fitness differences (48), revealed species-specific signatures that were associated with BGCs and their encoded specialized metabolites (Figure 1). Species-specific genomic patterns translated to metabolite production (Figure 3A) and demonstrated that specialized metabolism can be viewed as a phylogenetically conserved trait. Broadly, our results suggest that specialized metabolites contribute to functional differences capable of promoting ecological differentiation and subsequent fine-scale diversification in microbial communities.

Previous reports in soil bacteria have shown that the production of volatile organic compounds exhibit a clear taxonomic signal (49), suggesting bacterial specialized metabolites represent conserved phylogenetic traits resembling chemotaxonomy in plants and fungi (5, 50). Similarly, we observed a distinct phylogenetic signal at both the BGC and metabolite levels, indicating that vertical inheritance is a major driver of BGC evolution. While horizontal gene transfer (HGT) may play a major role in expanding BGC diversity, the prevailing view that BGCs are rapidly gained and lost (21-24) remains difficult to discern given that genomic signatures (e.g., GC% and tetranucleotide bias) can be lost over time (51). As such, the time scales in which these events occur are highly relevant when considering the processes driving BGC evolution. For example, within the genus *Streptomyces*, it was recently estimated that a single gene is acquired through HGT every 100k years (52) with the capacity to vertically maintain BGCs (i.e., the antibiotic streptomycin) for >40 MYA (53). By examining the distribution of BGCs among closely related *Salinispora* species, we also detected a strong signal of vertical inheritance, even among BGCs that likely originated from ancestral HGT events. This suggests that the collection of BGCs within *Salinispora* largely represents molecules that provide a fitness advantage.

While the BGCs are largely maintained by vertical descent, the evolutionary processes driving their genetic and, ultimately, chemical diversification remain poorly understood. We observed these processes affecting BGC evolution at multiple levels. First, at a broad level, vertical descent, BGC loss events, and rare HGT events account for the composition of *Salinispora* BGCs. High rates of homologous recombination between closely related strains combined with barriers to interspecies transfer (due to dispersal limitation and genetic/ecological divergence) maintain cohesion between phylogeny and BGC composition. Second, within a gene cluster family (GCF), interspecies differences in auxiliary gene composition affected compound production. This was observed in both the *sta* and *sal* BGCs (Figures 4 and S10A) where different species preferentially produced different staurosporine and salinosporamide analogs. Similarly, this pattern is attributed to high rates of homologous recombination of small sections of the BGC (Table 1), allowing for gain/loss of auxiliary genes while maintaining the core biosynthetic machinery (21). Finally, at the gene level, interspecies differences in nonsynonymous SNPs within the AT_1_ domain of the *salA* PKS gene were linked to the production of different analogs (Figure 4D). At this level, selection and the accumulation of genetic polymorphisms can further accelerate genetic and metabolic differentiation (54). Given that BGCs can be maintained over hundreds of millions of years via vertical descent, our results provide evidence that the evolutionary dynamics driving BGC evolution can diverge in association with speciation events. Collectively, these observations reveal that BGC evolution in *Salinispora* proceeds largely through gene gain/loss events and constrained recombination that contributes to interspecies diversity at the gene, pathway, and metabolite levels.

While genetic analyses offer insights to the evolutionary processes structuring BGC distributions, the resulting metabolite is ultimately the biological trait undergoing selection. Our observation that the collection of BGCs observed among 118 *Salinispora* genomes are largely structured by vertical inheritance suggests their products provide a selective advantage, although confirmation of their ecological roles is needed. While we did not observe strong evidence of recent selective sweeps (i.e., reduced genetic diversity (29)), specific metabolites were fixed at the species level. This is exemplified by the production of rifamycin S by *S. arenicola* (Figure 3B). The elevated genetic diversity observed in the *rif* BGC is neutral in terms of compound production, suggesting that negative selection has removed deleterious mutations that compromise the structure of the compound. In other cases, the diversity of analogs produced by a single BGC highlights the difficulty in viewing metabolic diversity as a single genotype to phenotype trait. Together, our observations are consistent with recently proposed models of BGC evolution (55, 56) where neutral (nondirectional) processes allow for exploration of metabolic diversity while negative selection removes deleterious compounds in favor of advantageous molecules. This relaxation of purifying selection might resemble phenotypic plasticity, in the form of chemical diversity, allowing for the exploration of ecological landscapes through beneficial natural products. In this manner, BGC plasticity could facilitate the exploitation of local environmental resources through population-specific pathway evolution as an initial step in species diversification.

There is a growing appreciation that metabolite-mediated interactions can influence evolutionary fitness landscapes (53, 55). Inversely, understanding these dynamics, especially from an evolutionary perspective, can facilitate future natural product discovery efforts. Here, we show that BGC diversification had direct consequences on compound production and analog distribution, supporting links between specialized metabolite and species diversification. While the rates at which BGCs are acquired and possibly lost via HGT remain unknown, current BGC distributions revealed species-specific signatures that are consistent with niche defining functional traits. These findings reinforce the concept that our knowledge of microbial diversity is critical for natural product discovery (57) and highlights the value of targeting rare or poorly studied taxa. Ultimately, resolving the roles of metabolite-mediated interactions as drivers of ecological differentiation will help inform our understanding of microbial diversity and community dynamics.

## MATERIALS AND METHODS

See Supplemental Information.

## SUPPLEMENTAL FIGURES AND TABLES

Figure S1. Abundance of BGCs across the bacterial domain.

Figure S2. Distribution of species-specific orthologous proteins.

Figure S3. *Salinispora* BGC network and BGC rank abundance curve

Figure S4. Pairwise ANI values for whole-genome comparisons and BGCs.

Figure S5. Molecular clock analysis and BGC phylogenies.

Figure S6. Recombination network based on gene flow in *Salinispora*.

Figure S7. Selection coefficients in conserved BGC genes.

Figure S8. Structures of *Salinispora* specialized metabolites.

Figure S9. Representative BGCs for the *sta* and *spo* BGCs.

Figure S10. Detailed BGC architecture of the salinosporamide BGC.

Figure S11. Molecular network for salinosporamide production.

Figure S12. Salinosporamide AT domain phylogeny

Figure S13. Signature motifs in the active site of the AT_1_ domain

Figure S14. LC/MS analysis for sporolide and salinilactam

Table S1. Characteristics of *Salinispora* strains.

Table S2. Reference information on nine BGCs.

Table S3. High-resolution MS for compounds associated with nine BGCs.

## DATA AVAILABILITY

All genomes are publicly available (Table S1). Public datasets for all metabolomic spectra files are available at massive.ucsd.edu (MSV000085890). All other data and relevant code used can be found at https://github.com/alex-b-chase/salBGCevol.

## Author Contributions

ABC and PRJ designed and conceptualized the research project. ABC, DS, and MNM performed culture and metabolomic experiments. DS, MNM, DGM analyzed metabolomicdata. ABC analyzed genomic data and conducted statistical analyses. ABC, DS, and PRJ wrote manuscript.

