## Supplemental Figures for "Vertical inheritance governs biosynthetic gene cluster evolution and chemical diversification"

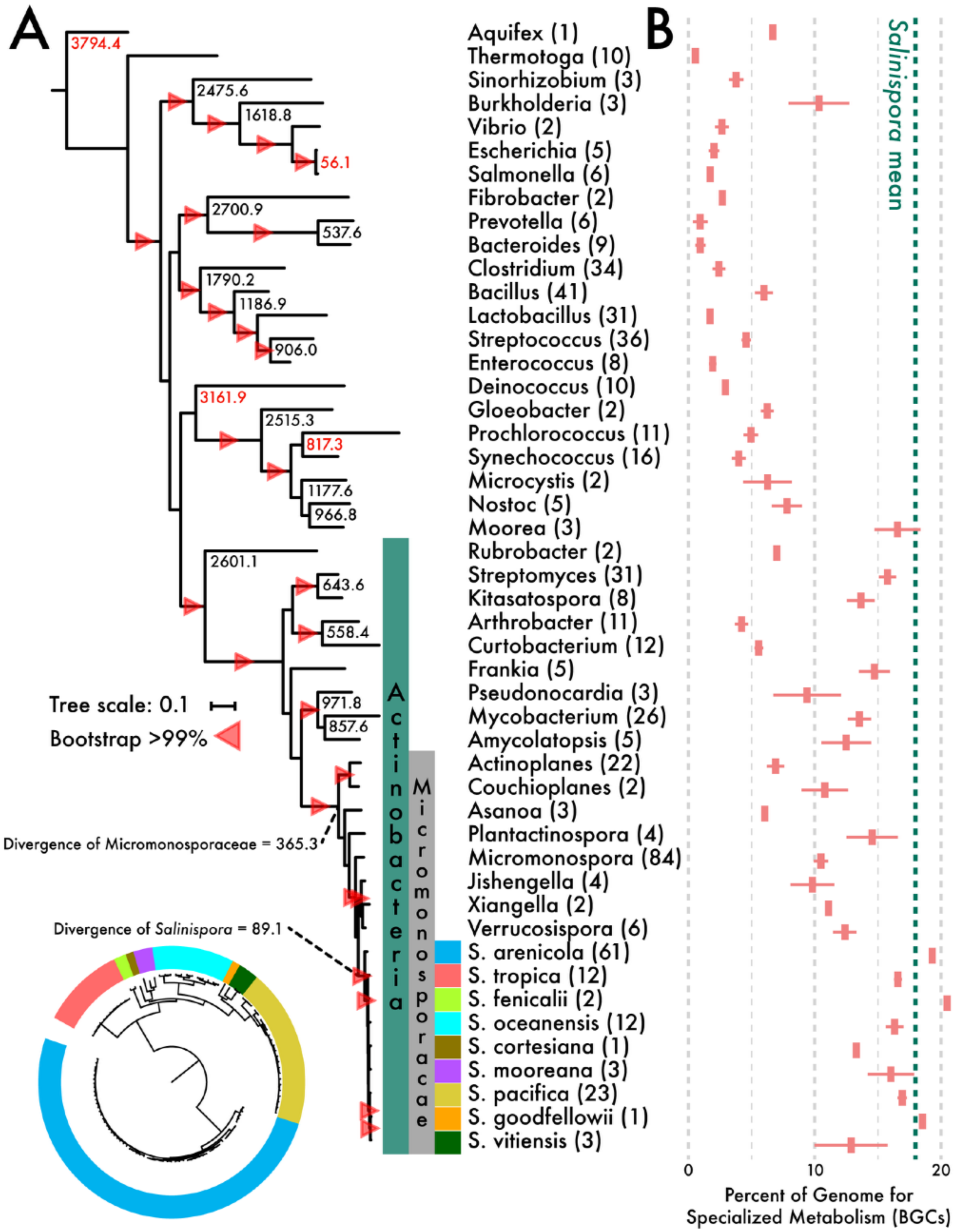


**Supplemental Figure S1.** Distribution of BGCs across the bacterial domain. **A)** Multilocus phylogenetic analysis of conserved, single-copy marker genes. Estimated time of divergences (millions of years; MYA) are denoted at nodes with reference calibration points in red. The phylum *Actinobacteria* and the family Micromonosporaceae are denoted for reference to *Salinispora*. **B)** Average percentage of genome dedicated to specialized metabolism based on annotated BGCs. Number of genomes analyzed per genus in parentheses.

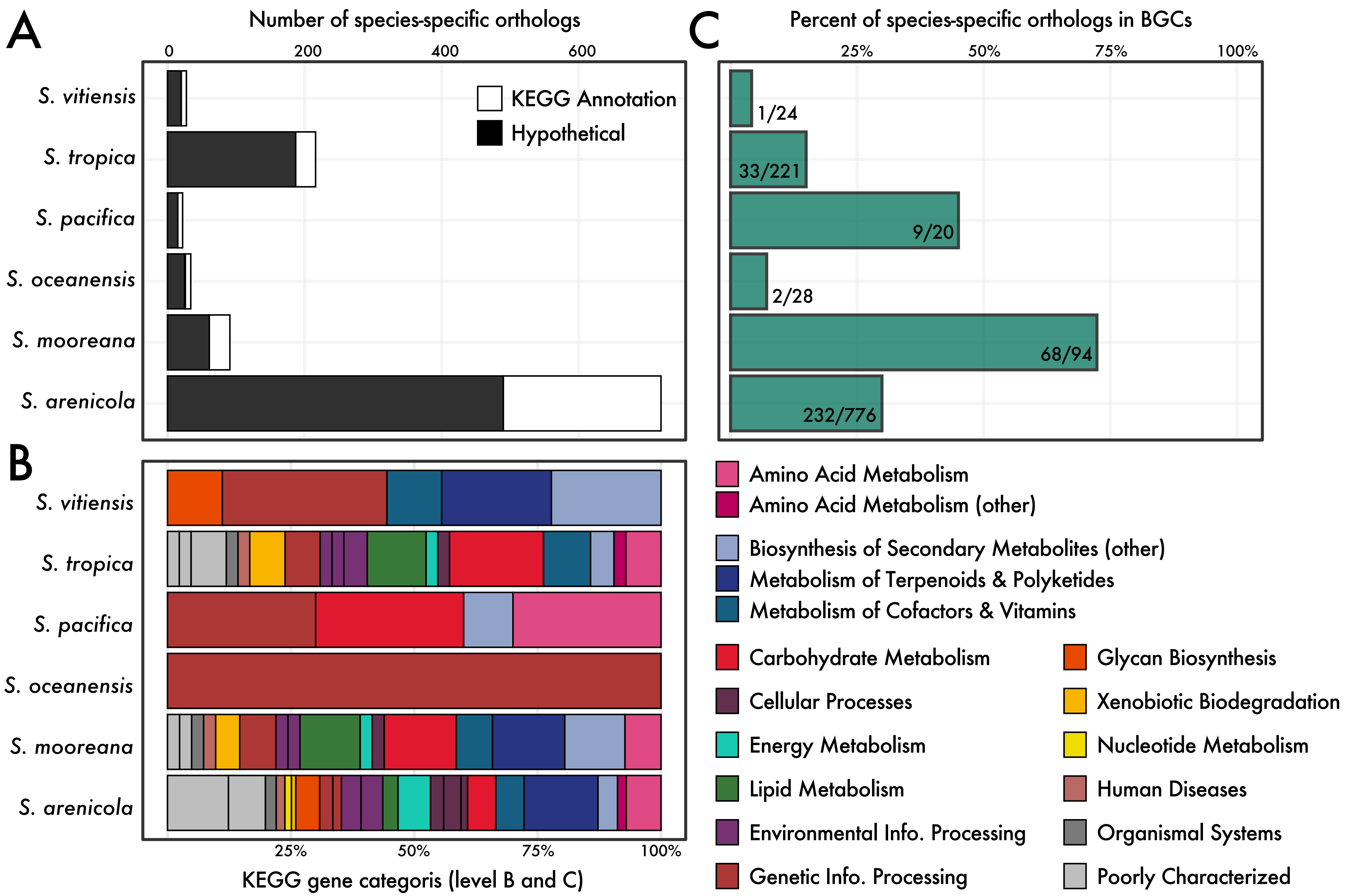


**Supplemental Figure S2.** Distribution of species-specific orthologous proteins in *Salinispora* species with ≥3 genomes per species. **A)** Number of hypothetical and annotated species-specific orthologous proteins. **B)** KEGG functional annotations for Panel A orthologs. **C)** Percent of total species-specific orthologs located within biosynthetic gene cluster (BGC) boundaries. Species only with ≥3 genomes were analyzed.

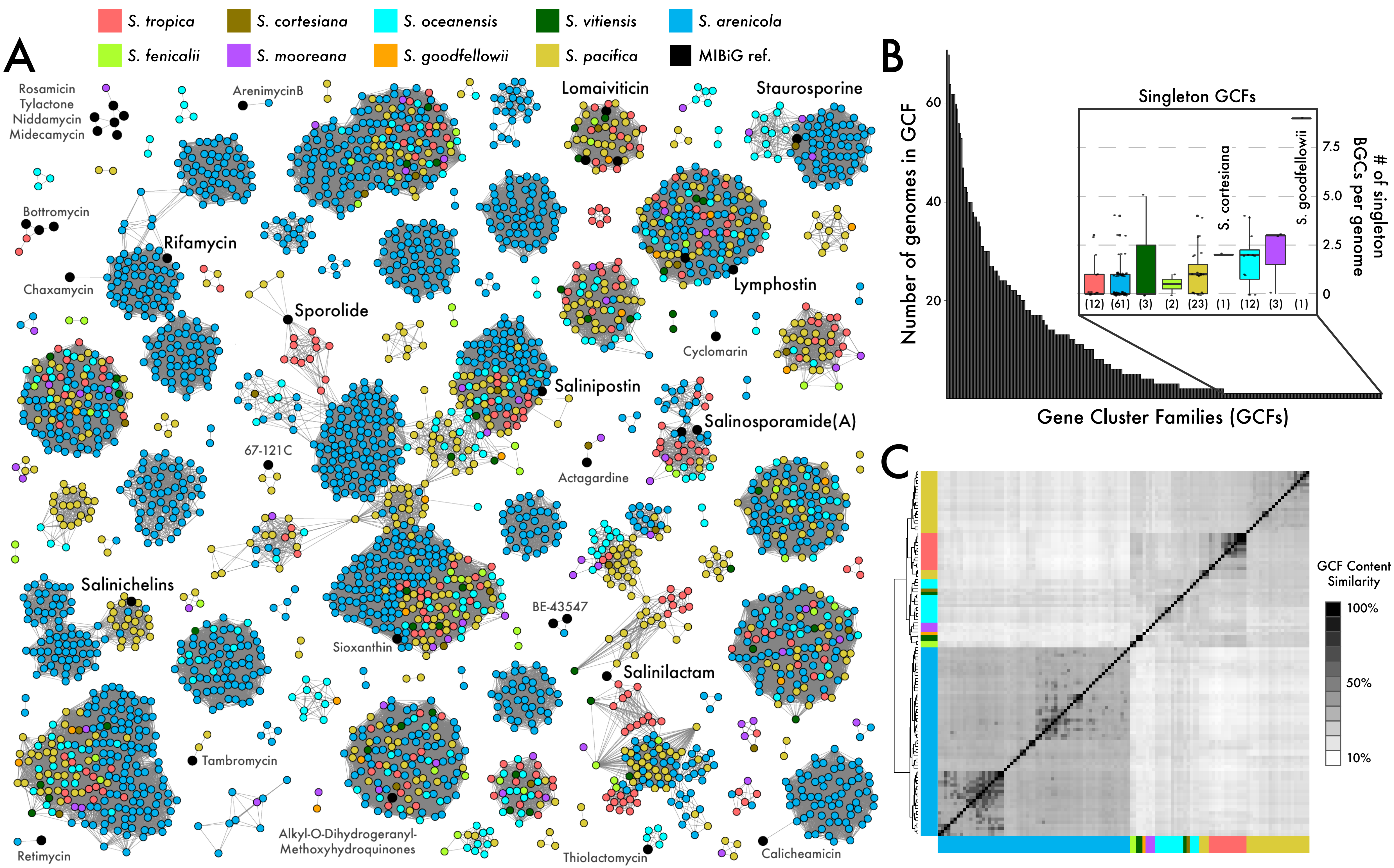


**Supplemental Figure S3.** Distribution of *Salinispora* biosynthetic gene clusters (BGCs). **A)** BGC network with gray edges representing squared similarity scores. Each node represents an individual BGC from a strain and is colored by species. Black nodes correspond to MIBiG reference BGC(s). **B)** Rank abundance curve of gene cluster families (GCFs). Inset shows average distribution of singleton GCF across genomes by species. **C)** BGC module similarity between strains. Heatmap was generated from a Jaccard distance matrix.

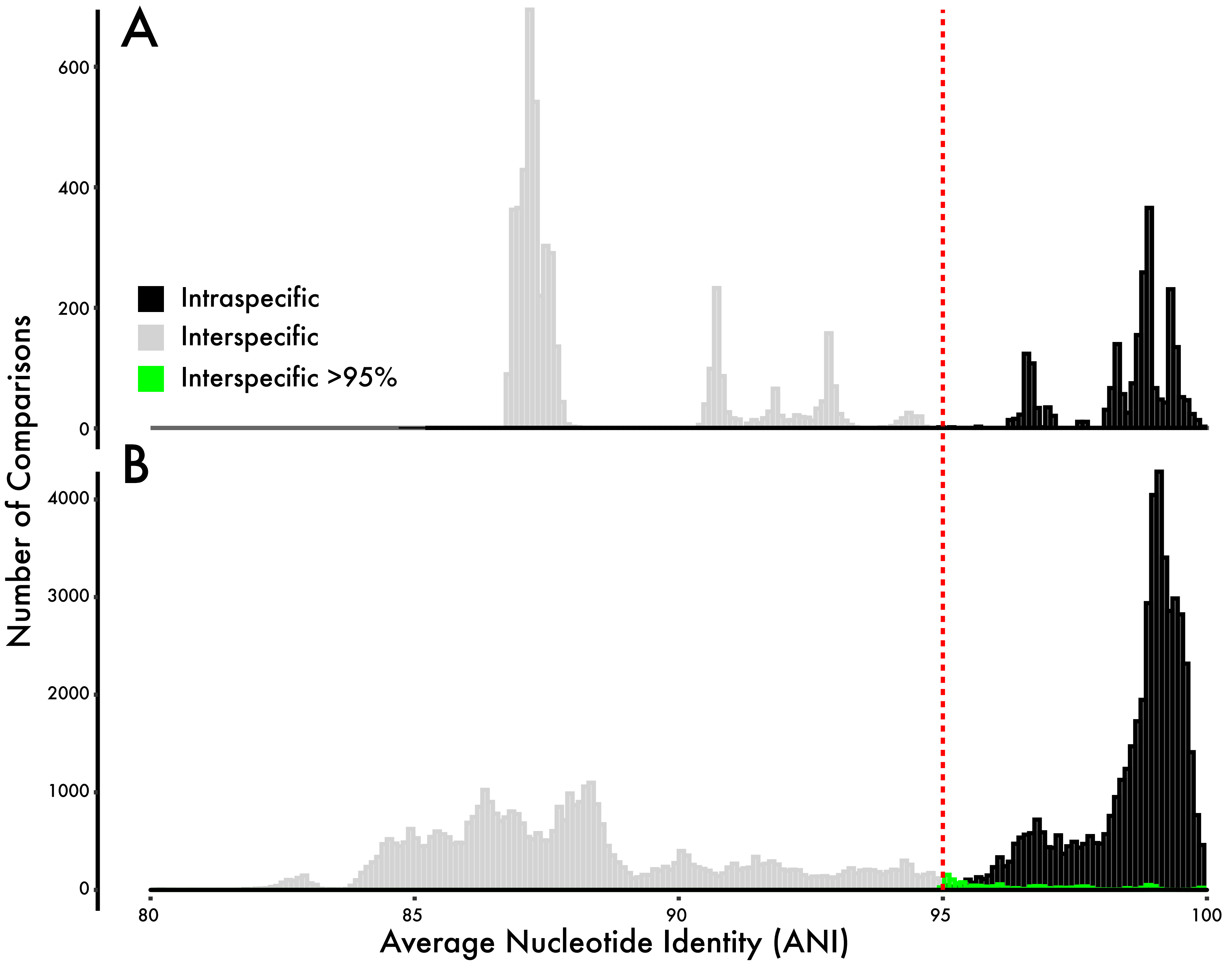


**Supplemental Figure S4.** Pairwise average nucleotide identity (ANI) values for **A)** whole-genome comparisons and **B)** biosynthetic gene clusters (BGCs). Colors indicate whether the comparison was between strains from the same species (black) or different species (gray). Interspecific BGC comparisons higher than the value used for species delineation (red dashed line, ANI ≥ 95%) are highlighted in green.

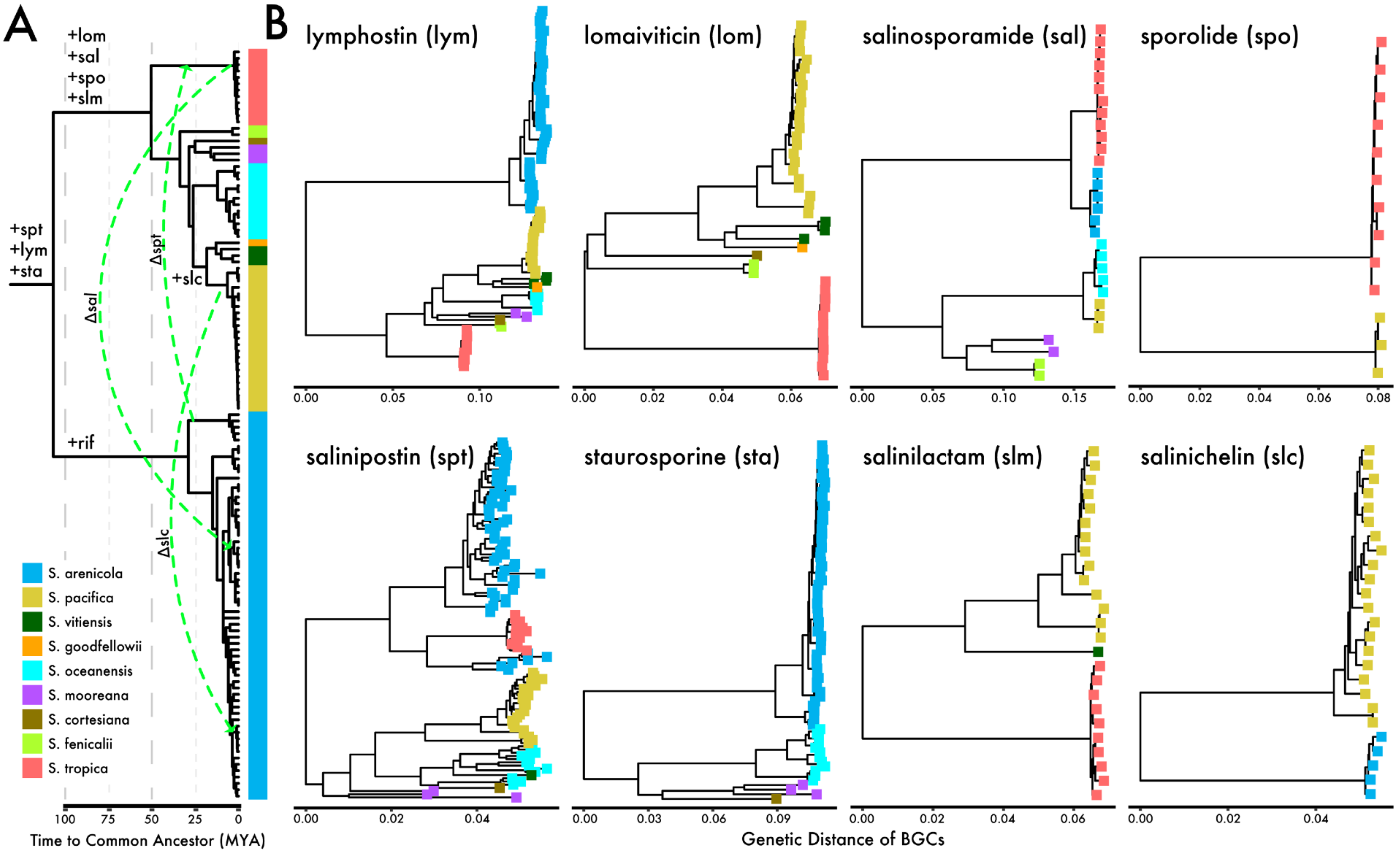


**Supplemental Figure S5.** Relationship between *Salinispora* species divergence and BGC diversification. **A)** *Salinispora* multilocus phylogeny with estimated time to common ancestor (millions of years; MYA). Reference BGCs are superimposed on phylogeny and refer to predicted acquisition events over the evolutionary history of *Salinispora*. **B)** Phylogenies of BGCs are congruent to the *Salinispora* core genome analysis except where denoted in panel A (green arrows) showing horizontal transfer of BGCs between species. Phylogenies generated from BGC nucleotide alignments (rifamycin not shown as it is only found in *S. arenicola*). Trees are midpoint rooted; scale bars represent genetic distance. Branch tips colored by species.

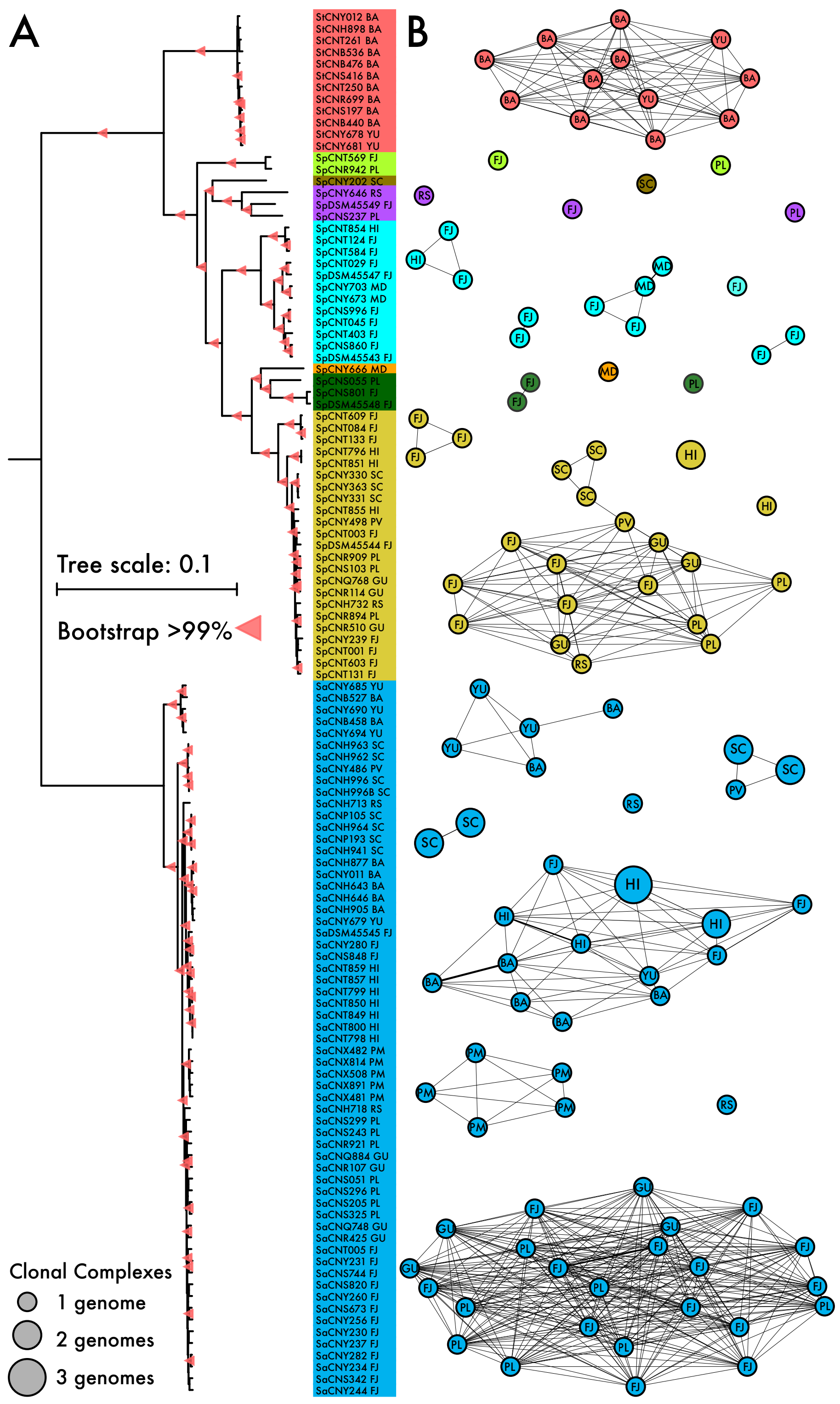


**Supplemental Figure S6.** Recombination network based on recent gene flow in *Salinispora* genomes. **A)** Core genome phylogeny (from Figure 1A). Colors denote species. Bar, 0.1 nucleotide substitutions per position. Bootstrap values indicate >99% support. **B)** Recombination network. Thicker edges represent increased recombination between strains. Nodes colored by species with site of isolation indicated (Supplementary Table S1). Node size indicates the number of clonal clusters (strains too similar to differentiate recombination and vertical inheritance).

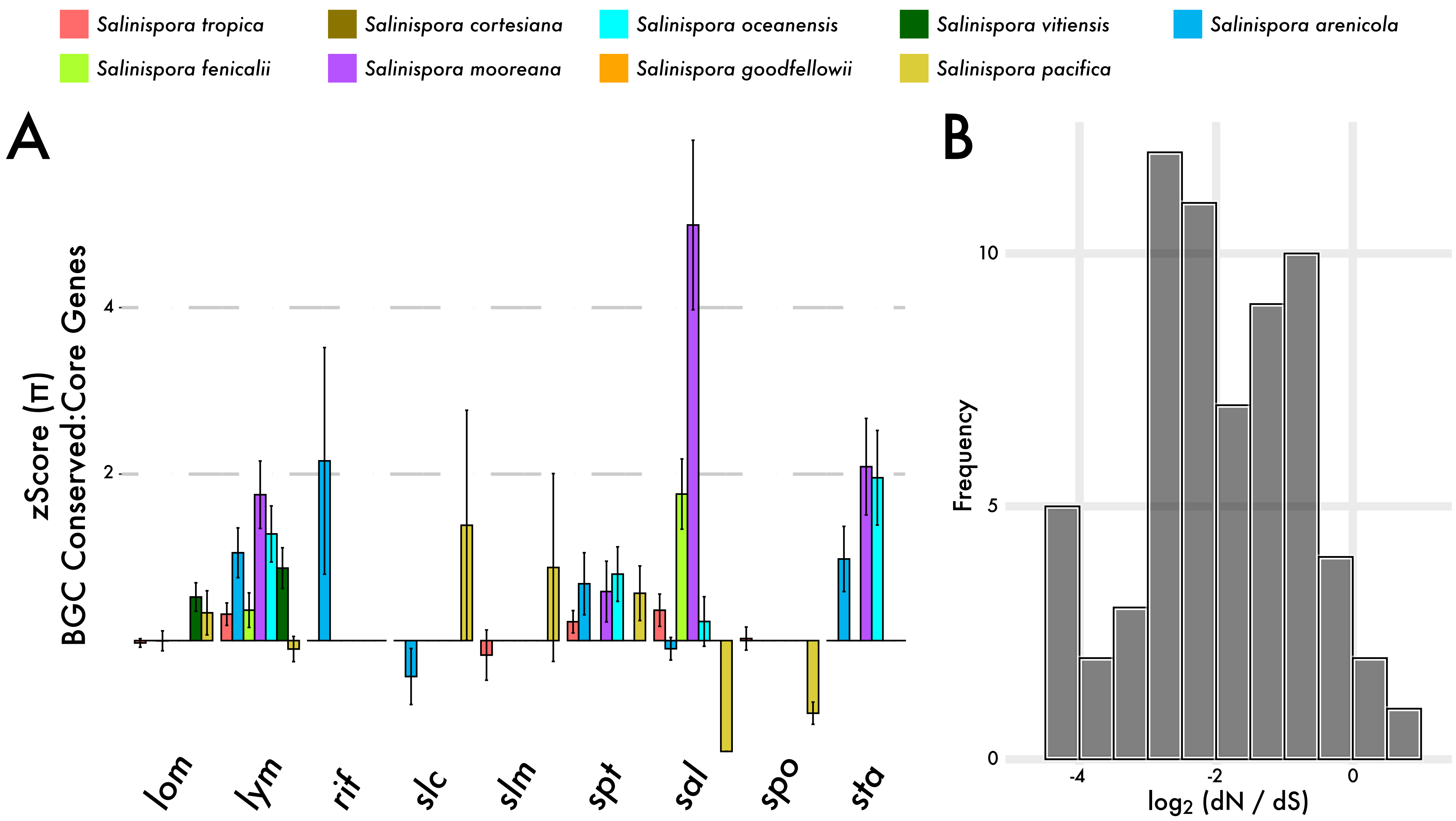


**Supplemental Figure S7.** Evidence for selection in conserved BGC genes. **A)** Nucleotide diversity (π) across nine BGCs. zScores reflect average π of conserved BGC genes relative to the core genome. **B)** Frequency of dN/dS ratios of conserved BGC genes.

### Salinichelins

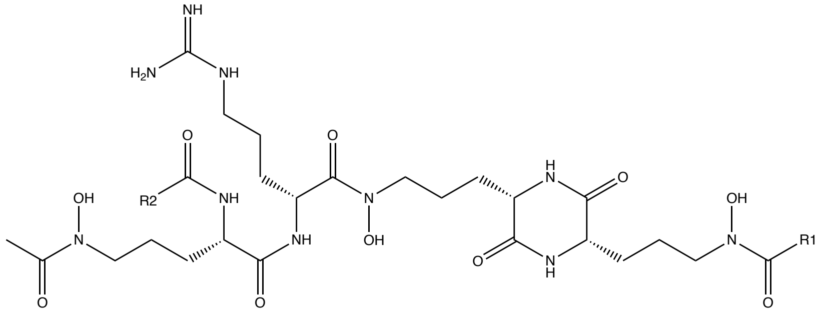

| | | |
|---|---|---|
| salinichelin A | R1 -H | R2 -Me |
| salinichelin B | R1 -Me | R2 -Me |
| salinichelin C | R1 -H | R2 -Et |

### Salinosporamides

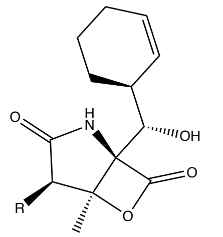

| | |
|---|---|
| salinosporamide A | R -Cl |
| salinosporamide B | R -Me |
| salinosporamide D | R -Et |
| salinosporamide E | R -Pr |
| salinosporamide K | R -H |

### Lomaiviticins

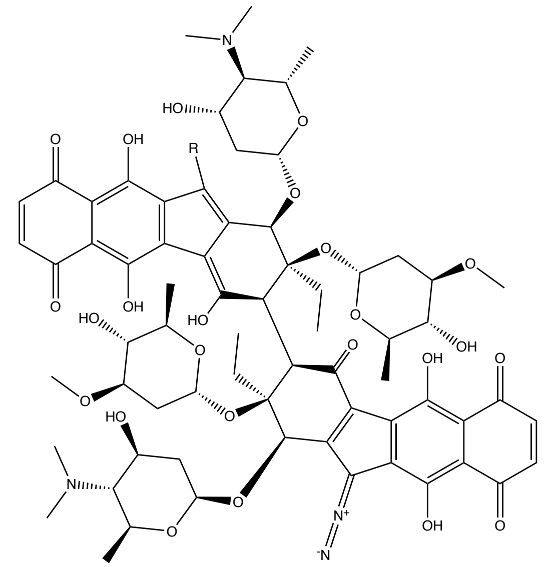

| | |
|---|---|
| lomaiviticin A | R $=N_2$ |
| lomaiviticin C | R -H |

### Sporolides

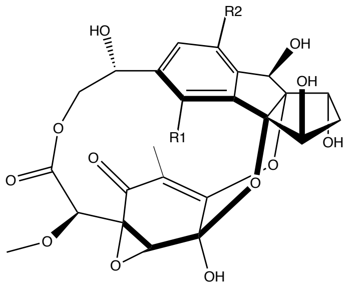

| | | |
|---|---|---|
| sporolide A | R1 -Cl | R2 -H |
| sporolide B | R1 -H | R2 -Cl |

### Salinilactams

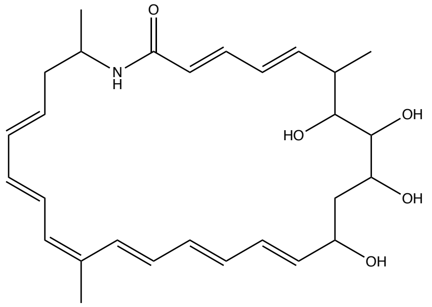

### Rifamycin S

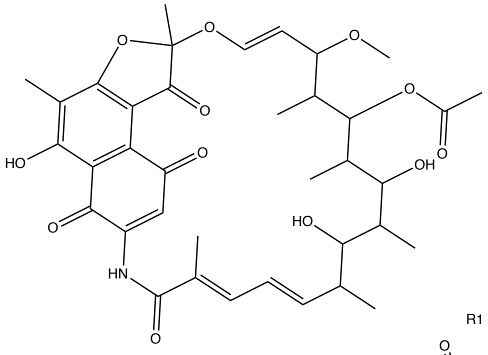

### Salinipostins

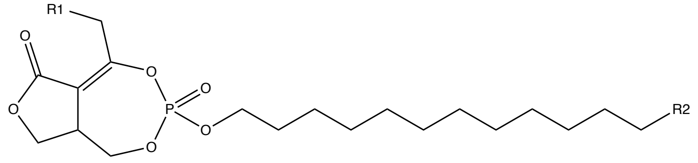

| | | |
|---|---|---|
| salinipostin A | R1 -Pr | R2 -Pr |
| salinipostin B | R1 -Pr | R2 -Et |
| salinipostin C | R1 -Pr | R2 -Me |
| salinipostin D | R1 -iso-Pr | R2 -Et |
| salinipostin E | R1 -iso-Pr | R2 -Me |
| salinipostin F | R1 -Et | R2 -Pr |
| salinipostin G | R1 -Et | R2 -Et |
| salinipostin H | R1 -Et | R2 -Me |
| salinipostin I | R1 -Me | R2 -Pr |
| salinipostin K | R1 -Me | R2 -Me |

### Staurosporines

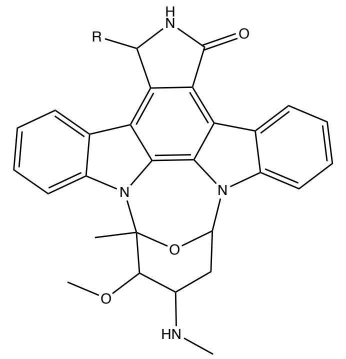

| | |
|---|---|
| staurosporine | R -H |
| 7-Hydroxystaurosporine | R -OH |

### Lymphostins

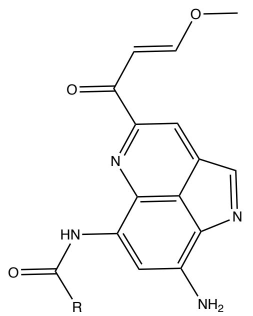

| | |
|---|---|
| lymphostin | R -Me |
| neolymphostin A | R -iso-Pr |
| neolymphostin B | R -Et |
| neolymphostin C | R -sec-Bu |
| neolymphostin D | R -iso-Bu |

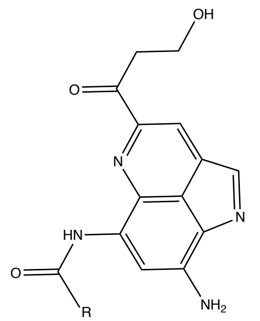

| | |
|---|---|
| lymphostinol | R -Me |
| neolymphostinol A | R -iso-Pr |
| neolymphostinol B | R -Et |
| neolymphostinol C | R -sec-Bu |
| neolymphostinol D | R -iso-Bu |

**Supplemental Figure S8.** Structures of nine *Salinispora* specialized metabolites and a subset of known analogs.

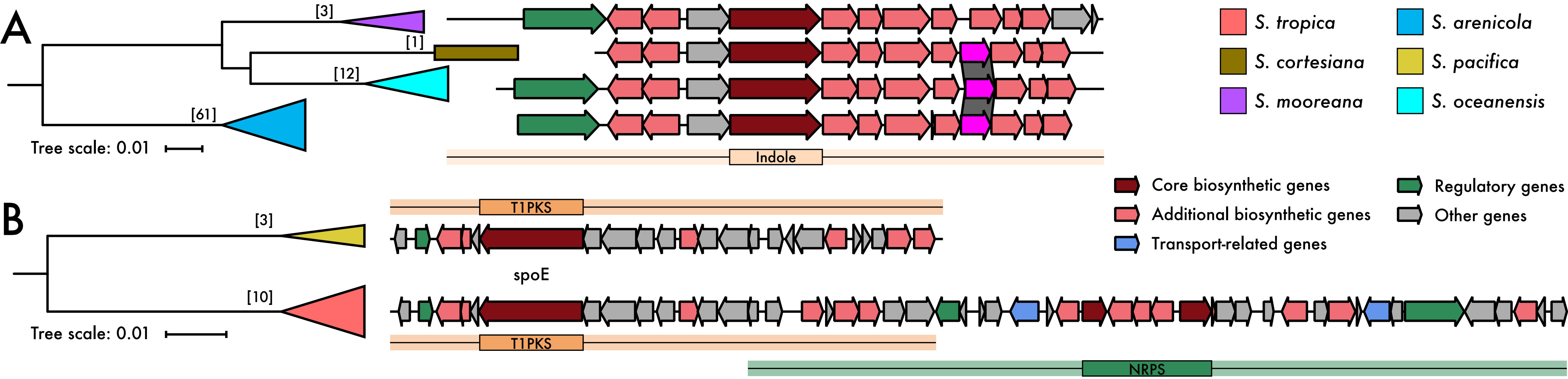


**Supplemental Figure S9.** Representative BGCs. **A)** *sta* BGC, encoding the staurosporines and **B)** *spo* BGC, encoding the sporolides. Phylogenies based on concatenated alignment of two essential BGC proteins (Table S2). Colors denote species. Bar, 0.01 amino acid substitutions per position. Brackets indicate number of genomes in each species encoding the BGC. Panel A also indicates the absence of the NAD-dependent dehydratase enzyme (magenta colored) in *S. mooreana*.

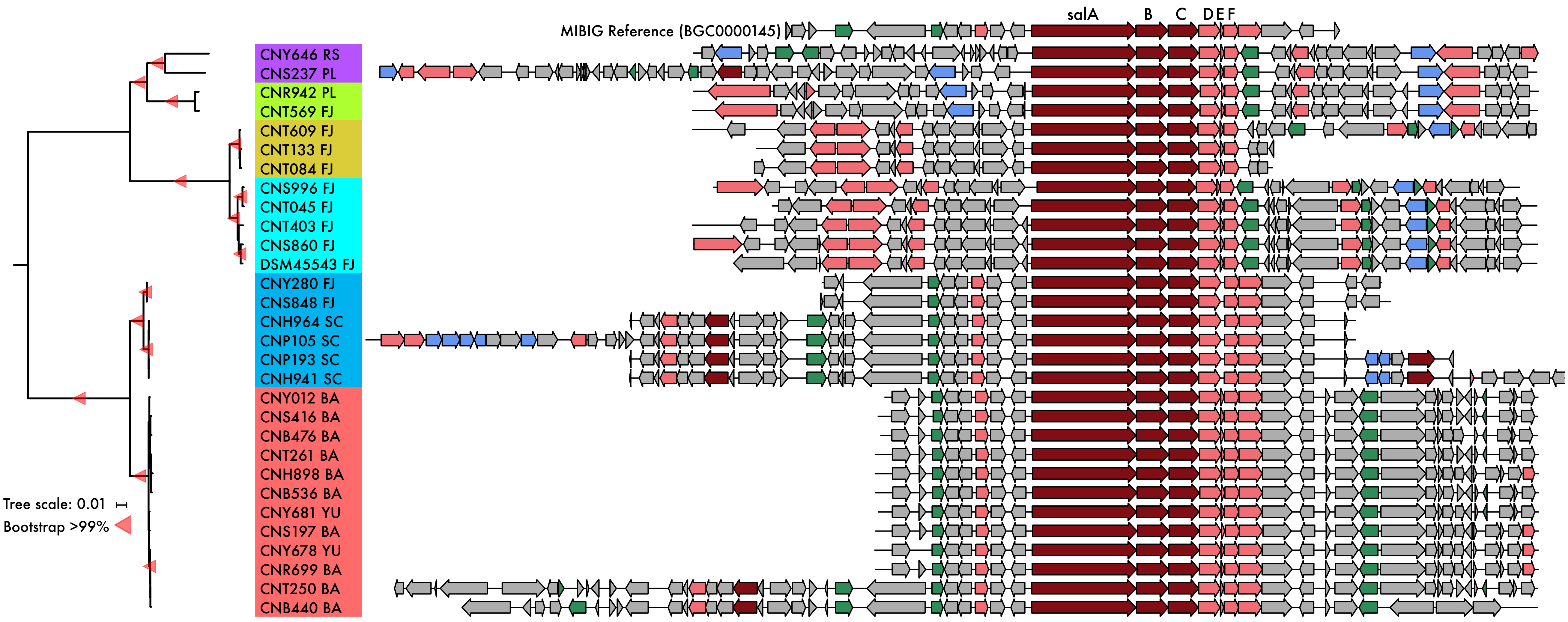


**Supplemental Figure S10.** Expanded version of Figure 4A-B showing the detailed BGC architecture across all strains encoding the salinosporamide BGC. The MIBIG *sal* BGC is shown for reference.

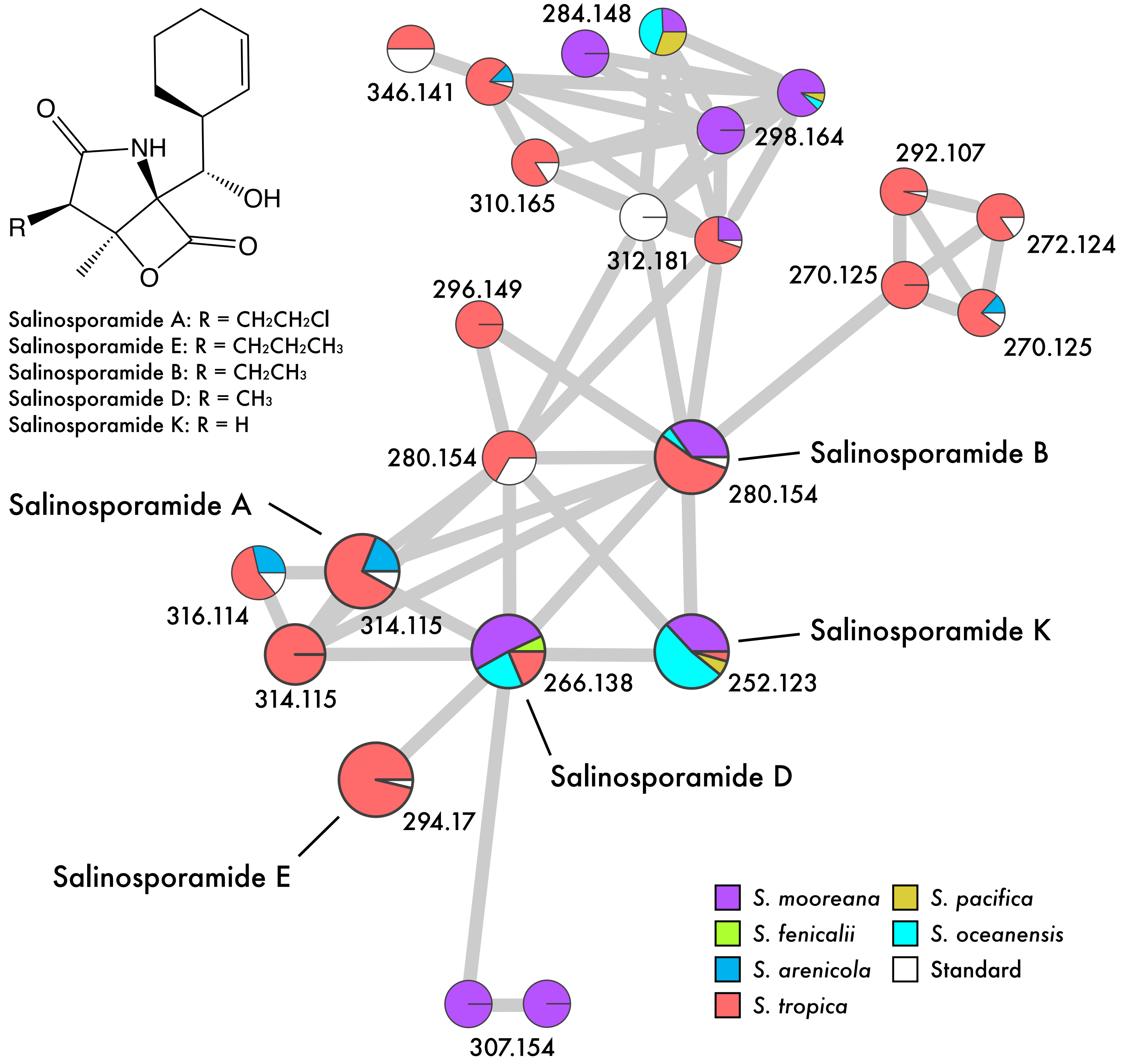


**Supplemental Figure S11.** Molecular network showing clustering of MS2 fragmentation spectra. Each node represents a unique fragmentation spectrum with the parent mass [M+H] denoted. Pie charts represent the number of MS2 scans detected for the parent mass, colored by species. Larger nodes depict known salinosporamides (labeled).

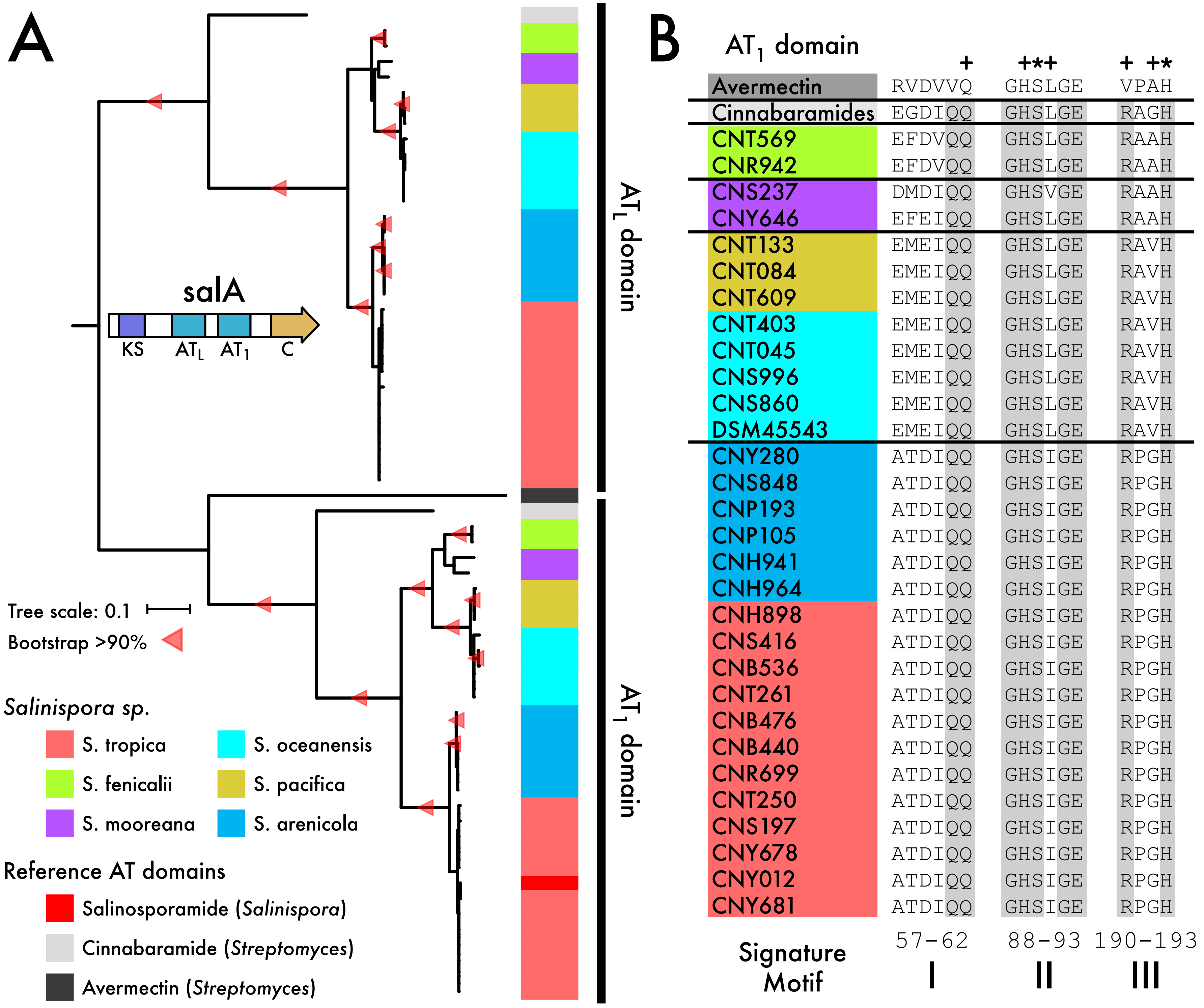


**Supplemental Figure S12.** Salinosporamide AT domain phylogeny and active site conservation. **A)** Phylogenetic analysis of the salinosporamide $AT_L$ (loading) and $AT_1$ (extension) domains with reference AT domains. **B)** Amino acid alignments for three (I-III) $AT_1$ domain signature motifs involved in substrate recognition. Predicted active site residues (+) and catalytic residues (*) from the avermectin AT domain indicated.

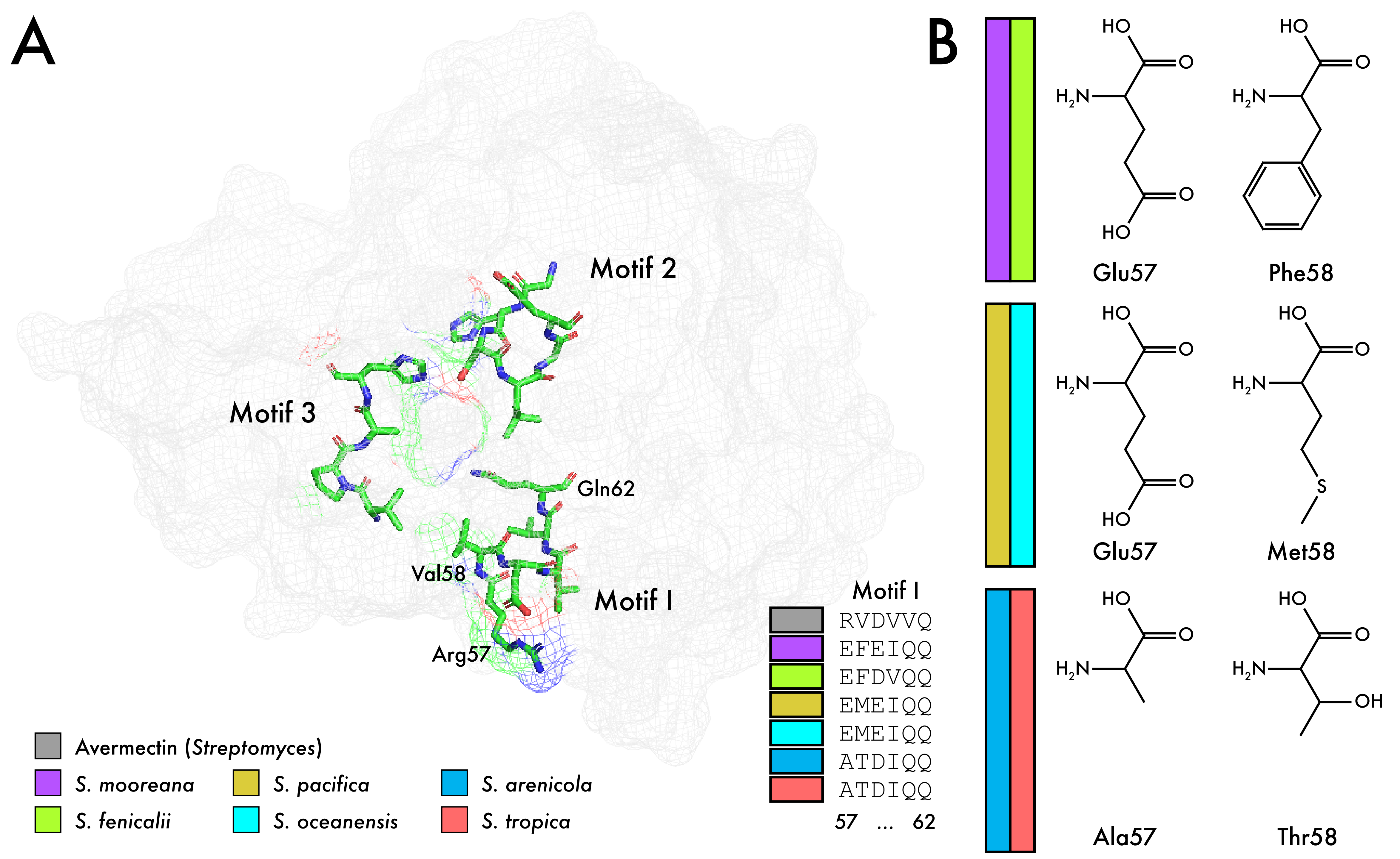


**Supplemental Figure S13.** Signature motifs in the active site of the AT domain. **A)** Mesh surface representation of the secondary structure protein model of the avermectin (PDB: 4RL1) AT domain. Amino acid residues in the three signature motifs involved in substrate recognition are colored using the green (carbon), blue (nitrogen), red (oxygen) format. **B)** Amino acid residues at positions 57-58 in signature motif I grouped by *Salinispora* species pairs.

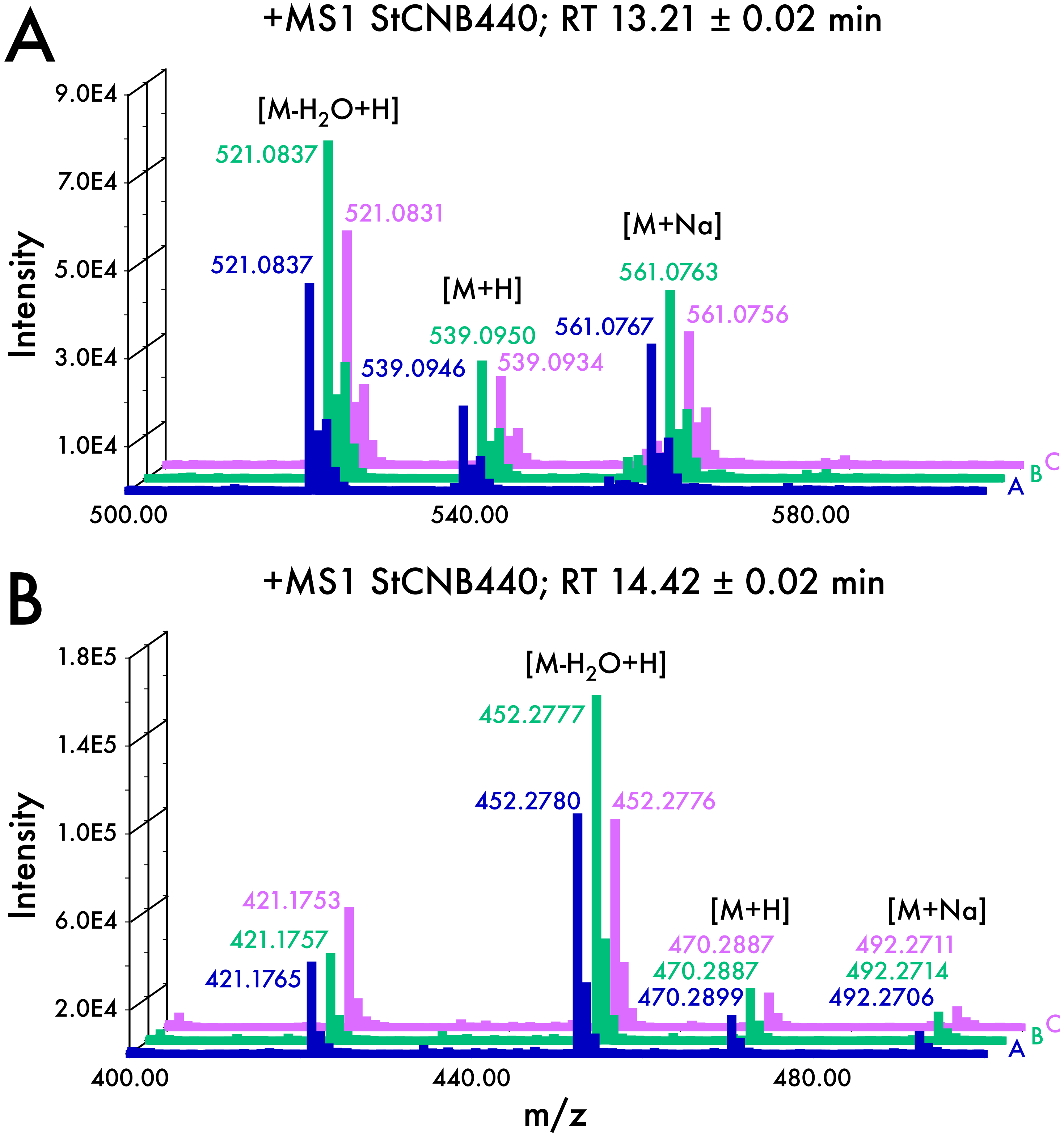


**Supplemental Figure S14.** LC/MS analysis for the identification of **A)** sporolide and **B)** salinilactam across three replicate grow-ups in *S. tropica* CNB440. MS1 analysis was used due to poor fragmentation of the parent ions in MS2 spectra.

| Strain ID | Species | Metadata | | | | | Genome Stats | | | | BGC Stats | |
|---|---|---|---|---|---|---|---|---|---|---|---|---|
| | | NCBI Taxon ID | Isolation Source | Location | Latitude-Longitude | Collection Year | Number of Contigs | Genome Length (bp) | Number of CDS | %GC | Number of BGC Fragments | Percent (%) of Genome for BGCs |
| CNB458 | arenicola | 1136425 | Marine Sediment | Bahamas | 26.626167, -77.919833 | 1989 | 132 | 5704144 | 5173 | 69 | 27 | 17.4% |
| CNB527 | arenicola | 1137250 | Marine Sediment | Bahamas | 22.350000, -74.016667 | 1989 | 120 | 5981449 | 5360 | 69 | 32 | 19.4% |
| CNH643 | arenicola | 1408326 | Marine Sediment | Bahamas | NA | 1999 | 85 | 5610201 | 5004 | 69 | 27 | 17.9% |
| CNH646 | arenicola | 1136427 | Marine Sediment | Bahamas | 26.600000, -77.883333 | 1999 | 58 | 5400492 | 4789 | 69 | 26 | 18.6% |
| CNH713 | arenicola | 1408327 | Marine Sediment | Red Sea | NA | 2000 | 43 | 5505299 | 4860 | 69 | 25 | 19.3% |
| CNH718 | arenicola | 1408328 | Marine Sediment | Red Sea | 27.566667, 33.930000 | 2000 | 54 | 5449370 | 4770 | 69 | 27 | 19.2% |
| CNH877 | arenicola | 1169176 | Marine Sediment | Bahamas | 24.018883, -74.544483 | 2000 | 93 | 5795920 | 5197 | 69 | 30 | 18.7% |
| CNH905 | arenicola | 1137251 | Marine Sediment | Bahamas | 24.018883, -74.544483 | 2000 | 105 | 5667031 | 5048 | 69 | 29 | 19.0% |
| CNH941 | arenicola | 1137252 | Marine Sediment | Sea of Cortez | NA | 2000 | 73 | 5761497 | 5218 | 69 | 29 | 19.3% |
| CNH962 | arenicola | 1169175 | Marine Sediment | Sea of Cortez | NA | 2000 | 87 | 5434989 | 4853 | 69 | 27 | 16.8% |
| CNH963 | arenicola | 1288085 | Marine Sediment | Sea of Cortez | NA | 2000 | 105 | 5440051 | 4869 | 69 | 27 | 17.0% |
| CNH964 | arenicola | 1136430 | Marine Sediment | Sea of Cortez | NA | 2000 | 70 | 5912993 | 5311 | 69 | 31 | 21.1% |
| CNH996 | arenicola | 168697 | Marine Sediment | Sea of Cortez | 24.824833, -110.586000 | 2001 | 151 | 5857846 | 5292 | 69 | 32 | 18.2% |
| CNH996B | arenicola | 1408330 | Marine Sediment | Sea of Cortez | 24.824833, -110.586000 | 2001 | 155 | 5939586 | 5367 | 69 | 33 | 17.8% |
| CNP105 | arenicola | 1169174 | Marine Sediment | Sea of Cortez | 24.856167, -110.588667 | 2001 | 69 | 5919675 | 5355 | 69 | 29 | 20.6% |
| CNP193 | arenicola | 1169170 | Marine Sediment | Sea of Cortez | 24.428500, -110.425333 | 2001 | 79 | 5766351 | 5223 | 69 | 29 | 19.4% |
| CNQ748 | arenicola | 1144929 | Marine Sediment | Guam | 13.365000, 144.646667 | 2002 | 90 | 5821375 | 5053 | 69 | 29 | 19.8% |
| CNQ884 | arenicola | 1408331 | Marine Sediment | Guam | 13.292417, 144.646550 | 2002 | 106 | 5838486 | 5153 | 69 | 31 | 19.1% |
| CNR107 | arenicola | 1169167 | Marine Sediment | Guam | 13.306033, 144.650433 | 2002 | 48 | 5767406 | 5090 | 69 | 28 | 19.8% |
| CNR425 | arenicola | 1298919 | Marine Sediment | Guam | 13.279567, 144.653750 | 2002 | 70 | 5705881 | 5072 | 69 | 29 | 19.2% |
| CNR921 | arenicola | 1136429 | Marine Sediment | Palau | 7.160583, 134.368783 | 2004 | 47 | 5458536 | 4875 | 69 | 25 | 18.5% |
| CNS051 | arenicola | 1169178 | Marine Sediment | Palau | 7.151500, 134.355667 | 2004 | 93 | 5853040 | 5169 | 69 | 29 | 17.7% |
| CNS205 | arenicola | 391037 | Marine Sediment | Palau | 7.108333, 134.253333 | 2004 | 1 | 5786361 | 5070 | 69 | 27 | 21.4% |
| CNS243 | arenicola | 1288087 | Marine Sediment | Palau | 7.295067, 134.515833 | 2004 | 70 | 5761551 | 5015 | 69 | 32 | 21.0% |
| CNS296 | arenicola | 1408332 | Marine Sediment | Palau | 7.170250, 134.352750 | 2004 | 47 | 5797291 | 5134 | 69 | 29 | 21.1% |
| CNS299 | arenicola | 1288086 | Marine Sediment | Palau | 7.240517, 134.380917 | 2004 | 63 | 5746562 | 5067 | 69 | 32 | 20.7% |
| CNS325 | arenicola | 1408333 | Marine Sediment | Palau | 7.326500, 134.436917 | 2004 | 59 | 5577349 | 4949 | 69 | 28 | 20.6% |
| CNS342 | arenicola | 1408334 | Marine Sediment | Fiji | NA | 2004 | 77 | 5586402 | 4951 | 69 | 27 | 18.6% |
| CNS673 | arenicola | 1144930 | Marine Sediment | Fiji | -18.761117, 178.563817 | 2006 | 83 | 5875699 | 5262 | 69 | 27 | 17.5% |
| CNS744 | arenicola | 1169168 | Marine Sediment | Fiji | -18.755700, 178.523750 | 2006 | 89 | 5741573 | 5133 | 69 | 29 | 20.8% |
| CNS820 | arenicola | 1408335 | Marine Sediment | Fiji | -18.713433, 178.490633 | 2006 | 66 | 5606043 | 4966 | 69 | 30 | 20.6% |
| CNS848 | arenicola | 1408351 | Marine Sediment | Fiji | -18.406233, 178.164250 | 2006 | 84 | 6036042 | 5374 | 69 | 31 | 20.7% |
| CNT005 | arenicola | 1137255 | Marine Sediment | Fiji | -18.780217, 178.550317 | 2006 | 67 | 5769669 | 5074 | 69 | 30 | 21.0% |
| CNT798 | arenicola | 1137256 | Marine Sediment | Hawaii | 20.634767, -156.493767 | 2008 | 66 | 5657349 | 5034 | 69 | 30 | 19.4% |
| CNT799 | arenicola | 1169172 | Marine Sediment | Hawaii | 20.634767, -156.493767 | 2008 | 58 | 5621489 | 4969 | 69 | 29 | 19.8% |
| CNT800 | arenicola | 1137253 | Marine Sediment | Hawaii | 20.634767, -156.493767 | 2008 | 51 | 5526283 | 4885 | 69 | 30 | 19.8% |
| CNT849 | arenicola | 1137264 | Marine Sediment | Hawaii | 20.634767, -156.493767 | 2008 | 51 | 5720436 | 5109 | 69 | 29 | 19.2% |
| CNT850 | arenicola | 1136428 | Marine Sediment | Hawaii | 20.634767, -156.493767 | 2008 | 67 | 5621922 | 4976 | 69 | 28 | 19.1% |
| CNT857 | arenicola | 1137254 | Marine Sediment | Hawaii | 20.639636, -156.450408 | 2008 | 92 | 5712025 | 5119 | 69 | 30 | 19.4% |
| CNT859 | arenicola | 1169163 | Marine Sediment | Hawaii | 20.639636, -156.450408 | 2008 | 74 | 5822487 | 5210 | 69 | 29 | 18.7% |
| CNX481 | arenicola | 1169169 | Marine Sediment | Palmyra | 5.869633, 162.077450 | 2009 | 73 | 5570975 | 4974 | 69 | 27 | 19.1% |
| CNX482 | arenicola | 1136426 | Marine Sediment | Palmyra | 5.892417, 162.076817 | 2009 | 89 | 5711575 | 5119 | 69 | 28 | 18.6% |
| CNX508 | arenicola | 1137257 | Marine Sediment | Palmyra | 5.870550, 162.124850 | 2009 | 96 | 5632853 | 5014 | 69 | 26 | 19.0% |
| CNX814 | arenicola | 1169171 | Marine Sediment | Palmyra | 5.869633, 162.077450 | 2009 | 65 | 5688460 | 5081 | 69 | 28 | 19.5% |
| CNX891 | arenicola | 1137258 | Marine Sediment | Palmyra | 5.869633, 162.077450 | 2009 | 92 | 5731186 | 5128 | 69 | 27 | 20.3% |
| CNY011 | arenicola | 1169162 | Marine Sediment | Bahamas | 26.563967, -77.890600 | 2010 | 82 | 5722126 | 5089 | 69 | 28 | 19.0% |
| CNY230 | arenicola | 1408336 | Sponge | Fiji | NA | 2011 | 95 | 6066259 | 5433 | 69 | 33 | 20.3% |
| CNY231 | arenicola | 1169179 | Marine Sediment | Fiji | -17.657700, 178.843467 | 2007 | 69 | 5832889 | 5151 | 69 | 28 | 19.8% |
| CNY234 | arenicola | 1169165 | Sponge | Fiji | NA | 2011 | 51 | 5569418 | 4928 | 69 | 28 | 19.2% |
| CNY237 | arenicola | 1169185 | Marine Sediment | Fiji | -17.273367, 177.100867 | 2008 | 67 | 5713643 | 5140 | 69 | 27 | 19.0% |
| CNY244 | arenicola | 1408337 | Marine Sediment | Fiji | -16.671833, -179.870833 | 2009 | 41 | 5497013 | 4863 | 69 | 28 | 19.7% |
| CNY256 | arenicola | 1169166 | Sponge | Fiji | NA | 2011 | 79 | 5781119 | 5169 | 69 | 26 | 17.9% |
| CNY260 | arenicola | 1169177 | Marine Sediment | Fiji | -17.996300, 179.188067 | 2007 | 63 | 5775769 | 5153 | 69 | 29 | 19.8% |
| CNY280 | arenicola | 1169173 | Marine Sediment | Fiji | -18.019483, 179.235633 | 2007 | 89 | 6066372 | 5445 | 69 | 30 | 21.0% |
| CNY282 | arenicola | 1169164 | Sponge | Fiji | NA | 2011 | 67 | 5755139 | 5137 | 69 | 27 | 18.6% |
| CNY486 | arenicola | 1408340 | Marine Sediment | Puerto Vallarta | 20.544783, -105.289667 | 2011 | 113 | 5680170 | 5112 | 69 | 31 | 17.5% |
| CNY679 | arenicola | 1408342 | Marine Sediment | Yucatan | 24.669417, -82.908483 | 2012 | 76 | 5760922 | 5180 | 69 | 26 | 17.7% |
| CNY685 | arenicola | 1408343 | Marine Sediment | Yucatan | 18.406217, -87.412517 | 2012 | 192 | 6269423 | 5796 | 69 | 35 | 18.2% |
| CNY690 | arenicola | 1408344 | Marine Sediment | Yucatan | 18.564333, -87.421250 | 2012 | 155 | 5877395 | 5346 | 69 | 27 | 17.4% |
| CNY694 | arenicola | 1408345 | Marine Sediment | Yucatan | 20.324650, -87.027117 | 2012 | 109 | 5792452 | 5245 | 69 | 30 | 18.7% |
| DSM45545 | arenicola | 999546 | Marine Sediment | Fiji | -18.761117, 178.563817 | 2006 | 1 | 5867409 | 5156 | 69 | 26 | 22.5% |
| CNY202 | cortesiana | 1305843 | Marine Sediment | Sea of Cortez | 25.950322, -111.306283 | 2008 | 204 | 5183058 | 4815 | 69 | 23 | 13.3% |
| CNR942 | fenicalii | 1169187 | Marine Sediment | Palau | 7.266667, 134.466667 | 2004 | 52 | 5473304 | 4963 | 69 | 24 | 20.5% |
| CNT569 | fenicalii | 1137263 | Marine Sediment | Fiji | -18.254333, 178.085000 | 2008 | 39 | 5234686 | 4732 | 69 | 22 | 20.4% |
| CNY666 | goodfellowii | 1408341 | Marine Sediment | Madeira Islands | 32.648350, -16.822750 | 2012 | 106 | 5726457 | 5204 | 70 | 27 | 18.5% |
| CNS237 | mooreana | 1288089 | Marine Sediment | Palau | 7.355200, 134.440150 | 2004 | 67 | 5243134 | 4735 | 69 | 23 | 17.9% |
| CNY646 | mooreana | 1408356 | Sponge | Red Sea | NA | 2006 | 73 | 5210670 | 4772 | 69 | 25 | 17.9% |
| DSM45549 | mooreana | 999545 | Marine Sediment | Fiji | -18.421683, 178.140883 | 2006 | 3 | 5198196 | 4883 | 69 | 15 | 12.3% |
| CNS860 | oceanensis | 1169186 | Marine Sediment | Fiji | -18.713433, 178.490633 | 2006 | 62 | 5357469 | 4918 | 69 | 22 | 16.0% |
| CNS996 | oceanensis | 1169289 | Marine Sediment | Fiji | -18.761117, 178.563817 | 2006 | 113 | 5661283 | 5174 | 69 | 27 | 18.3% |
| CNT029 | oceanensis | 1136418 | Marine Sediment | Fiji | -18.410433, 178.158233 | 2006 | 37 | 5236748 | 4743 | 69 | 19 | 13.5% |
| CNT045 | oceanensis | 1169190 | Marine Sediment | Fiji | -18.761117, 178.563817 | 2006 | 80 | 5771968 | 5270 | 69 | 28 | 18.6% |
| CNT124 | oceanensis | 1169188 | Marine Sediment | Fiji | -18.780217, 178.550317 | 2006 | 47 | 5164130 | 4691 | 69 | 20 | 17.5% |
| CNT403 | oceanensis | 1408353 | Marine Sediment | Fiji | -16.947117, 177.400333 | 2008 | 100 | 5394694 | 4940 | 69 | 23 | 16.2% |
| CNT584 | oceanensis | 1169191 | Marine Sediment | Fiji | -18.254333, 178.085000 | 2008 | 79 | 5217717 | 4782 | 69 | 23 | 17.4% |
| CNT854 | oceanensis | 1137265 | Marine Sediment | Hawaii | 20.637115, -156.450245 | 2008 | 104 | 5315422 | 4874 | 69 | 20 | 14.2% |
| CNY673 | oceanensis | 1408357 | Marine Sediment | Madeira Islands | 33.052583, -16.278333 | 2012 | 198 | 5799079 | 5321 | 69 | 22 | 13.5% |
| CNY703 | oceanensis | 1408346 | Marine Sediment | Madeira Islands | 32.536550, -16.532850 | 2012 | 135 | 5393461 | 4960 | 69 | 17 | 12.2% |
| DSM45543 | oceanensis | 999542 | Marine Sediment | Fiji | -18.399100, 178.034767 | 2006 | 1 | 5458289 | 5010 | 69 | 25 | 18.7% |
| DSM45547 | oceanensis | 1050199 | Marine Sediment | Fiji | -18.761117, 178.563817 | 2006 | 2 | 5447211 | 4864 | 69 | 22 | 19.7% |
| CNH732 | pacifica | 1408347 | Marine Sediment | Red Sea | 24.375833, 35.383833 | 2000 | 68 | 5188493 | 4684 | 69 | 21 | 17.9% |
| CNQ768 | pacifica | 1169193 | Marine Sediment | Guam | 13.518317, 144.790667 | 2002 | 91 | 5480836 | 4981 | 69 | 24 | 17.9% |
| CNR114 | pacifica | 1137260 | Marine Sediment | Guam | 13.286817, 144.649050 | 2002 | 99 | 5866896 | 5371 | 69 | 23 | 15.6% |
| CNR510 | pacifica | 1408348 | Marine Sediment | Guam | 13.253171, 144.658865 | 2002 | 104 | 5715131 | 5249 | 69 | 26 | 16.6% |
| CNR894 | pacifica | 1137261 | Marine Sediment | Palau | 7.298233, 134.504200 | 2004 | 115 | 5518023 | 5001 | 69 | 22 | 16.7% |
| CNR909 | pacifica | 1408349 | Marine Sediment | Palau | 7.298233, 134.504200 | 2004 | 112 | 5455852 | 4958 | 69 | 26 | 17.9% |
| CNS103 | pacifica | 1137262 | Marine Sediment | Palau | 7.301317, 134.224150 | 2004 | 122 | 5584369 | 5091 | 69 | 23 | 15.5% |
| CNT001 | pacifica | 1136416 | Marine Sediment | Fiji | -18.743367, 178.541700 | 2006 | 82 | 5492313 | 4961 | 69 | 25 | 16.9% |
| CNT003 | pacifica | 1136417 | Marine Sediment | Fiji | -18.773783, 178.546400 | 2006 | 136 | 5469257 | 4933 | 69 | 28 | 16.7% |
| CNT084 | pacifica | 1136419 | Marine Sediment | Fiji | -18.406000, 178.018150 | 2006 | 138 | 5471270 | 5108 | 69 | 28 | 14.6% |
| CNT131 | pacifica | 1136420 | Marine Sediment | Fiji | -18.713433, 178.490633 | 2006 | 113 | 5527068 | 4909 | 69 | 29 | 19.2% |
| CNT133 | pacifica | 1408352 | Marine Sediment | Fiji | -18.713433, 178.490633 | 2006 | 126 | 5553074 | 5152 | 69 | 28 | 14.3% |
| CNT603 | pacifica | 1136424 | Marine Sediment | Fiji | -18.254333, 178.085000 | 2008 | 77 | 5395536 | 4866 | 69 | 22 | 17.6% |
| CNT609 | pacifica | 1169184 | Marine Sediment | Fiji | -17.259850, 177.107967 | 2008 | 95 | 5328928 | 4967 | 69 | 26 | 14.4% |
| CNT796 | pacifica | 1206102 | Marine Sediment | Hawaii | 20.634767, -156.493767 | 2008 | 99 | 5073739 | 4616 | 69 | 22 | 16.5% |
| CNT851 | pacifica | 1169180 | Marine Sediment | Hawaii | 20.634767, -156.493767 | 2008 | 100 | 5263032 | 4798 | 69 | 24 | 16.2% |
| CNT855 | pacifica | 1137266 | Marine Sediment | Hawaii | 20.634767, -156.493767 | 2008 | 85 | 5208629 | 4694 | 69 | 22 | 15.4% |
| CNY239 | pacifica | 1288090 | Marine Sediment | Fiji | -17.738500, -179.342167 | 2008 | 88 | 5368602 | 4823 | 69 | 25 | 16.9% |
| CNY330 | pacifica | 1169192 | Marine Sediment | Sea of Cortez | 28.989950, -113.398717 | 2008 | 138 | 5498479 | 4982 | 69 | 25 | 17.0% |
| CNY331 | pacifica | 1169181 | Marine Sediment | Sea of Cortez | 28.953183, -113.430867 | 2008 | 170 | 5740370 | 5184 | 69 | 28 | 18.0% |
| CNY363 | pacifica | 1408354 | Marine Sediment | Sea of Cortez | 28.953183, -113.430867 | 2008 | 211 | 5879613 | 5328 | 69 | 29 | 17.4% |
| CNY498 | pacifica | 1408355 | Marine Sediment | Puerto Vallarta | 20.701650, -105.564800 | 2011 | 74 | 5455855 | 4949 | 69 | 22 | 17.6% |
| DSM45544 | pacifica | 999543 | Marine Sediment | Fiji | -18.723567, 178.490383 | 2006 | 4 | 5464752 | 4926 | 69 | 22 | 22.3% |
| CNB440 | tropica | 369723 | Marine Sediment | Bahamas | 25.405212, -77.881818 | 1989 | 1 | 5183331 | 4662 | 69 | 17 | 18.2% |
| CNB476 | tropica | 1137247 | Marine Sediment | Bahamas | 26.466700, -77.083300 | 1989 | 100 | 5239804 | 4766 | 69 | 16 | 15.0% |
| CNB536 | tropica | 1136431 | Marine Sediment | Bahamas | 22.350000, -74.016700 | 1989 | 122 | 5552560 | 5101 | 69 | 25 | 16.0% |
| CNH898 | tropica | 1137248 | Marine Sediment | Bahamas | 24.544883, -75.928733 | 2000 | 114 | 5532979 | 5054 | 69 | 27 | 18.4% |
| CNR699 | tropica | 1169198 | Marine Sediment | Bahamas | 26.558783, -77.884933 | 2003 | 124 | 5608799 | 5097 | 69 | 21 | 16.2% |
| CNS197 | tropica | 1137249 | Marine Sediment | Bahamas | 26.571467, -77.886783 | 2004 | 75 | 5236719 | 4767 | 69 | 19 | 16.3% |
| CNS416 | tropica | 1169195 | Marine Sediment | Bahamas | 25.827633, -77.880833 | 2004 | 95 | 5180643 | 4704 | 69 | 18 | 15.8% |
| CNT250 | tropica | 1169196 | Marine Sediment | Bahamas | NA | 2007 | 59 | 5170212 | 4686 | 69 | 19 | 16.9% |
| CNT261 | tropica | 1288084 | Marine Sediment | Bahamas | NA | 2007 | 96 | 5510329 | 5031 | 69 | 23 | 17.8% |
| CNY012 | tropica | 1169197 | Marine Sediment | Bahamas | 26.563967, -77.890600 | 2010 | 154 | 5369729 | 4921 | 69 | 21 | 15.5% |
| CNY678 | tropica | 1408358 | Marine Sediment | Yucatan | 21.171033, -86.730200 | 2012 | 76 | 5107079 | 4639 | 69 | 18 | 15.9% |
| CNY681 | tropica | 1408359 | Marine Sediment | Yucatan | 20.324650, -87.027117 | 2012 | 56 | 4996884 | 4511 | 69 | 19 | 17.1% |
| CNS055 | vitiensis | 1169182 | Marine Sediment | Palau | 7.284867, 134.230417 | 2004 | 62 | 5263298 | 4746 | 70 | 24 | 18.7% |
| CNS801 | vitiensis | 1408350 | Marine Sediment | Fiji | -18.713433, 178.490633 | 2006 | 39 | 4969155 | 4587 | 69 | 12 | 9.9% |
| DSM45548 | vitiensis | 999544 | Marine Sediment | Fiji | -18.785850, 178.552583 | 2006 | 2 | 5097790 | 4738 | 69 | 12 | 10.1% |

**Supplemental Table S1.** Genomic and geographic characteristics of *Salinispora* strains.

| BGC | BGC abbreviation | BGC class | MIBiG reference(s) | MIBiG reference strain(s) | core biosynthetic genes | reference CDS | annotation of core gene | reference |
|---|---|---|---|---|---|---|---|---|
| salinosporamide(A) | sal | T1PKS-NRPS hybrid | BGC0000145.1 | S. tropica CNB-440 (strop) | salA | strop1024 | malonyl CoA-acyl carrier protein transacylase | Eustaquio et al. PNAS. 2009 |
| | | | BGC0001041.1 | | salB | strop1023 | AMP-dependent synthetase and ligase | |
| lymphostin | lym | T1PKS-NRPS hybrid | BGC0001007.1 | S. arenicola CNS-205 (sare) | lymA | sare3282 | beta-ketoacyl synthase | Miyanaga et al. JACS. 2011 |
| | | | BGC0001006.1 | S. tropica CNB-440 | lymB | sare3281 | methyltransferase | |
| lomaiviticin | lom | T2PKS | BGC0000241.1 | S. tropica CNB-440 | kinA (kinamycin homolog) | strop2223 | beta-ketoacyl synthase | Kersten et al. CBC. 2013 |
| | | | BGC0000240.1 | S. pacifica | kinB (kinamycin homolog) | strop2224 | beta-ketoacyl synthase | |
| rifamycin | rif | T1PKS | BGC0000137.1 | S. arenicola CNS-205 | rifE | sare1250 | beta-ketoacyl synthase | |
| | | | | | rifF | sare1251 | N-acetyltransferase | |
| staurosporine | sta | Indole | BGC0000827.1 | S. arenicola CNS-205 | vioB (violacein homolog) | sare2330 | dichlorochromopyrrolate synthase | Amos et al. PNAS. 2017 |
| | | | | | P450 | sare2331 | cytochrome P450 | |
| salinichelins | slc | NRPS | BGC0001767.1 | S. pacifica CNY331 | slcE | CNY331_02495 | AMP-dependent synthetase and ligase | Bruns et al. ISME. 2018 |
| | | | | | slcF | CNY331_02496 | mbtH-like protein | |
| sporolide | spo | T1PKS | BGC0000150.1 | S. tropica CNB-440 | spoE | strop2697 | beta-ketoacyl synthase | McGlinchey et al. JACS. 2008 |
| | | | | | spoE7 | strop2694 | cytochrome P450 | |
| salinilactam | slm | T1PKS | BGC0000142.1 | S. tropica CNB-440 | slmN | strop2768 | beta-ketoacyl synthase | Udwary et al. PNAS. 2007 |
| | | | | | TetR | strop2766 | TetR family transcriptional regulator | |
| salinipostin | spt | Butyrolactone | BGC0001458.1 | S. tropica CNB-440 | spt8 | strop4150 | F420-dependent glucose-6-phosphate dehydrogenase | Amos et al. PNAS. 2017 |
| | | | | | spt9 | strop4151 | gamma-butyrolactone synthase | |

**Supplemental Table S2.** Reference information on nine BGCs and their essential biosynthetic genes.

| Theoretical mass of compound | | | | | Feature-based molecular MS1 network (FBMN) | | | | GNPS MS2 Network | | | |
|---|---|---|---|---|---|---|---|---|---|---|---|---|
| chemical.formula | theoretical.mz.[M] | theoretical.[M+2H]2+ | theoretical.[M+H] | theoretical.[M+Na] | FBMN.feature | FBMN.M+H | FBMN.retentiontime | FBMN.deltaPPM | GNPS.M+H | GNPS.retentiontime | verification | notes |
| C68H80N6O24 | 1364.5224 | 683.269 | 1365.5302 | 1387.511618 | below MS1 detection | NA | NA | NA | 1365.53 | 11.0455 | GNPS hit | |
| C54H56N6O18 | 1076.36511 | 539.190355 | 1077.37291 | 1099.354328 | not found | NA | NA | NA | NA | NA | NA | |
| C68H82N4O24 | 1338.5319 | 670.27375 | 1339.5397 | 1361.521118 | feat122 | 670.2734 | 11.55433 | 0.59677 | 1339.54 | 11.5925 | GNPS hit | |
| C69H84N4O24 | 1352.54755 | 677.281575 | 1353.55535 | 1375.536768 | feat2375 | 677.2809 | 12.04354 | 1.03354 | 1353.55 | 12.0823 | GNPS network | |
| C70H86N4O24 | 1366.5632 | 684.2894 | 1367.571 | 1389.552418 | feat2837 | 684.2886 | 12.86878 | 1.16910 | 1367.57 | 12.9154 | GNPS network | |
| C16H14N4O3 | 310.10659 | N/A | 311.11439 | 333.095808 | feat2330 | 311.1132 | 10.6823 | 3.82496 | 311.113 | 10.5469 | GNPS hit | |
| C15H14N4O3 | 298.10659 | N/A | 299.11439 | 321.095808 | feat3548 | 299.1135 | 10.405796 | 2.97545 | 299.113 | 10.3854 | GNPS network | |
| C18H18N4O3 | 338.13789 | N/A | 339.14569 | 361.127108 | feat280 | 339.1447 | 12.29269 | 2.91910 | 339.1455 | 12.3242 | standard | |
| C17H16N4O3 | 324.12224 | N/A | 325.13004 | 347.111458 | feat317 | 325.1294 | 11.52677 | 1.96844 | 325.1295 | 11.5494 | GNPS network | |
| C19H20N4O3 | 352.15354 | N/A | 353.16154 | 375.142758 | feat651 | 353.1606 | 12.99634 | 2.66167 | 353.1603333 | 13.0299 | GNPS network | |
| C19H20N4O3 | 352.15354 | N/A | 353.16134 | 375.142758 | not found | NA | NA | NA | NA | NA | NA | |
| C17H18N4O3 | 326.13789 | N/A | 327.14569 | 349.127108 | feat2498 | 327.1446 | 11.86884 | 3.33185 | 327.1446667 | 11.8418 | GNPS network | |
| C16H16N4O3 | 312.12224 | N/A | 313.13004 | 335.111458 | feat288 | 313.1301 | 11.060237 | 0.19161 | 313.1296667 | 11.0506 | GNPS network | |
| C18H20N4O3 | 340.15354 | N/A | 341.16134 | 363.142758 | not found | NA | NA | NA | NA | NA | NA | |
| C18H20N4O3 | 340.15354 | N/A | 341.16134 | 363.142758 | not found | NA | NA | NA | NA | NA | NA | |
| C36H45NO12 | 683.29418 | N/A | 684.30198 | 706.283398 | not found | NA | NA | NA | NA | NA | NA | |
| C35H45NO10 | 639.30435 | N/A | 640.31215 | 662.293568 | feat2446 | 640.3083 | 16.18047 | 6.01269 | 640.31 | 16.1759 | GNPS hit | |
| C39H49NO14 | 755.31531 | N/A | 756.32311 | 778.304528 | not found | NA | NA | NA | NA | NA | NA | |
| C37H45NO12 | 695.29418 | N/A | 696.30198 | 718.283398 | feat1897 | 696.3013 | 20.83004 | 0.97659 | 696.301 | 20.4245 | GNPS hit | |
| C37H45NO12 | 695.29418 | N/A | 696.30198 | 718.283398 | feat1895 | 718.2828 | 20.83297 | 0.83254 | 718.283 | 20.7864 | GNPS hit | |
| C37H47NO12 | 697.30983 | N/A | 698.31763 | 720.299048 | not found | NA | NA | NA | NA | NA | NA | |
| C35H45NO11 | 655.29926 | N/A | 656.30706 | 678.288478 | not found | NA | NA | NA | 656.306 | 14.8977 | standard | |
| C15H20ClNO4 | 313.1081 | N/A | 314.1159 | 336.097318 | feat444 | 314.1158 | 16.36164 | 0.31835 | 314.115 | 16.2339 | standard | |
| C15H21NO4 | 279.1471 | N/A | 280.1549 | 302.136318 | feat192 | 280.1546 | 15.29792 | 1.07084 | 280.154 | 15.2867 | standard | |
| C14H18ClNO3 | 283.0975 | N/A | 284.1053 | 306.086718 | not found | NA | NA | NA | NA | NA | NA | |
| C14H19NO4 | 265.1314 | N/A | 266.1392 | 288.120618 | feat1777 | 266.1387 | 14.07203 | 1.87872 | 266.138 | 13.9311 | GNPS network | |
| C16H23NO4 | 293.1627 | N/A | 294.1705 | 316.151918 | feat475 | 294.17 | 16.50833 | 1.69969 | 294.17 | 16.5396 | GNPS network | |
| C15H20ClNO4 | 313.1081 | N/A | 314.1159 | 336.097318 | not found | NA | NA | NA | NA | NA | NA | epimeric of salA |
| C14H19NO4 | 265.1314 | N/A | 266.1392 | 288.120618 | feat1812 | 266.1387 | 13.79697 | 1.87872 | NA | NA | NA | epimeric of salD |
| C16H23NO4 | 293.1627 | N/A | 294.1705 | 316.151918 | not found | NA | NA | NA | NA | NA | NA | epimeric of salE |
| C16H22ClNO4 | 327.1237 | N/A | 328.1315 | 350.112918 | not found | NA | NA | NA | NA | NA | NA | |
| C15H20ClNO3 | 297.1132 | N/A | 298.121 | 320.102418 | not found | NA | NA | NA | NA | NA | NA | |
| C13H17NO4 | 251.1158 | N/A | 252.1236 | 274.105018 | feat1203 | 252.1233 | 13.01222 | 1.18989 | 252.123 | 13.0249 | GNPS network | |
| C26H46N10O10 | 658.33984 | N/A | 659.34764 | 681.329058 | not found | NA | NA | NA | NA | NA | NA | |
| C27H48N10O10 | 672.35549 | N/A | 673.36329 | 695.344708 | not found | NA | NA | NA | NA | NA | NA | |
| C27H48N10O10 | 672.35549 | N/A | 673.36329 | 695.344708 | not found | NA | NA | NA | NA | NA | NA | |
| C28H50N10O10 | 686.37114 | N/A | 687.37894 | 709.360358 | not found | NA | NA | NA | NA | NA | NA | |
| C28H50N10O10 | 686.37114 | N/A | 687.37894 | 709.360358 | not found | NA | NA | NA | NA | NA | NA | |
| C29H52N10O10 | 700.38679 | N/A | 701.39459 | 723.376008 | not found | NA | NA | NA | NA | NA | NA | |
| C29H52N10O10 | 700.38679 | N/A | 701.39459 | 723.376008 | not found | NA | NA | NA | NA | NA | NA | |
| C28H39NO5 | 469.28282 | N/A | 470.29062 | 492.272038 | feat1513 | 470.2892 | 14.40072 | 3.01941 | 470.289 | 14.4235 | manual MS1 analysis | |
| C24H23ClO12 | 538.0878 | N/A | 539.0956 | 561.077018 | feat1798 | 539.0931 | 13.19771 | 4.63740 | 539.094 | 12.8363 | manual MS1 analysis | isomers |
| C24H23ClO12 | 538.0878 | N/A | 539.0956 | 561.077018 | feat1820 | 539.0946 | 12.21168 | 1.85496 | 539.094 | 12.8363 | manual MS1 analysis | isomers |
| C25H45O6P | 472.29538 | N/A | 473.30318 | 495.284598 | not found | NA | NA | NA | NA | NA | standard | |
| C24H43O6P | 458.27973 | N/A | 459.28753 | 481.268948 | not found | NA | NA | NA | NA | NA | NA | |
| C23H41O6P | 444.26408 | N/A | 445.27188 | 467.253298 | not found | NA | NA | NA | NA | NA | NA | |
| C24H43O6P | 458.27973 | N/A | 459.28753 | 481.268948 | not found | NA | NA | NA | NA | NA | NA | |
| C23H41O6P | 444.26408 | N/A | 445.27188 | 467.253298 | not found | NA | NA | NA | NA | NA | NA | |
| C24H43O6P | 458.27973 | N/A | 459.28753 | 481.268948 | not found | NA | NA | NA | NA | NA | NA | |
| C23H41O6P | 444.26408 | N/A | 445.27188 | 467.253298 | not found | NA | NA | NA | NA | NA | NA | |
| C22H39O6P | 430.24843 | N/A | 431.25623 | 453.237648 | not found | NA | NA | NA | NA | NA | standard | |
| C23H41O6P | 444.26408 | N/A | 445.27188 | 467.253298 | not found | NA | NA | NA | NA | NA | NA | |
| C22H39O6P | 430.24843 | N/A | 431.25623 | 453.237648 | not found | NA | NA | NA | NA | NA | NA | |
| C21H37O6P | 416.23278 | N/A | 417.24058 | 439.221998 | feat1727 | 417.2381 | 18.79558 | 5.94381 | 417.238 | 18.0170 | negative MS1 analysis | |
| C28H26N4O4 | 482.19541 | N/A | 483.20321 | 505.184628 | feat1615 | 483.2021 | 11.57084 | 2.29717 | 483.202 | 12.0621 | GNPS hit | |
| C28H26N4O3 | 466.20049 | N/A | 467.20829 | 489.189708 | feat169 | 467.2073 | 12.19044 | 2.11897 | 467.207 | 12.8656 | standard | |
| C28H24N4O5 | 496.17467 | N/A | 497.18247 | 519.163888 | feat2460 | 497.1842 | 13.0138 | 3.47961 | 497.1835 | 13.0631 | GNPS network | |
| C28H27N4O3+ | 467.20777 | N/A | 468.21557 | 490.196988 | not found | NA | NA | NA | NA | NA | NA | |

**Supplemental Table S3.** High-resolution MS, retention time, and predicted chemical formula for associated compounds produced by nine BGCs.
