## Supplemental Information for "Vertical inheritance governs biosynthetic gene cluster evolution and chemical diversification"

**MATERIALS & METHODS**

**Phylogenomic and flexible genome analyses**

We downloaded 118 *Salinispora* genomes representing strains isolated from globally-distributed marine sediments (96% of isolates) and sponges (13). Previously, we assigned each strain to one of nine species based on genotypic and phenotypic characteristics (14) (Supplementary Table 1). For each genome, protein-coding regions and gene annotations were assigned using Prokka v1.13.3 (58) and orthologs shared across all genomes were identified with Roary v3.12.0 (59) based on a minimum sequence identity of 85%. The resulting 2106 potential orthologs were individually aligned using Clustal Omega (60) and screened for complete codon reading frames. The final 2011 single-copy orthologs were used to create a concatenated core genome alignment and reconstructed phylogenetic relatedness using RAxML v.8.2.10 (61) under the general time reversal model with a gamma distribution for 100 replicates. Any orthologs not shared among all strains were assigned to the flexible genome. Species-specific flexible genes (defined as genes shared by all strains within a species but not observed in any other species) were assigned functional annotation with GhostKOALA against the nonredundant set of KEGG genes (62) in all *Salinispora* species where we had adequate representation with ≥3 genomes per species.

For molecular clock analyses, we selected a representative genome from 40 genera spanning the bacterial domain (see below for more information on genera) including a representative genome from the nine *Salinispora* species. The representative genomes were used to build a MLST phylogeny with 50 single-copy marker genes. Briefly, protein sequences of the ribosomal marker genes were extracted with HMMer (63), aligned with Clustal Omega, and concatenated for phylogenetic analyses with RAxML under the WAG protein model for 100 replicates. The resulting alignment and phylogeny were used to calculate divergence times with a maximum likelihood clock in the RelTime (64) tool in MEGAX (65). Calibration points were set for the evolution of Cyanobacteria (2500-3500 MYA), divergence of *Salmonella* and *Escherichia* (50-150 MYA), divergence of *Prochlorococcus* and *Synechococcus* (644-823 MYA), and the origin of bacteria (3500-3800 MYA), as previously described (66). Using the divergence times of *Salinispora*, we recalibrated the molecular clock analyses with all 118 *Salinispora* genomes under identical parameters with *Micromonospora* as the outgroup.

**BGC identification and network visualization**

Each *Salinispora* genome was analyzed with antiSMASH v5.0-beta (67) to predict biosynthetic gene clusters (BGCs) associated with specialized metabolism. Identified BGCs were then analyzed with BiG-SCAPE v1.0.0 (68) to determine relationships with experimentally characterized BGCs in the MIBiG database (69) and group BGCs into gene cluster families (GCFs). The resulting GCFs were clustered based on a combination of protein domain sequence similarity and shared protein domains (distance <0.3). Pairwise distances (squared similarity scores ranging from 0-1) based on the BiG-SCAPE output were visualized in a network with a ForceAtlas layout in Gephi v0.9.2 (https://gephi.org/). The resulting network revealed that some GCFs were highly similar to one another as evident by them grouping into larger clusters (e.g., lomaiviticin cluster composed of two GCFs). Therefore, we the network connectivity to provide a second, higher level of BGC clustering by grouping nodes within the network into modules (set at a 1.0 resolution). The resulting modularity (M=0.927) detected highly connected modules within the network that collapsed similar GCFs into 206 BGC modules.

To address the relative importance of geographic location and phylogenetic relatedness on *Salinispora* genetic diversity, we computed a Jaccard distance matrix using both flexible genome content and GCF composition. For the flexible genome, the distance matrix was used to generate a neighbor-joining tree based on 1000 re-samplings and visualized with a heatmap showing gene content similarity across strains. For GCF composition, the distance matrices for both GCFs and BGC modules were used to construct non-metric multidimensional scales (NMDS) ordination plots. No major statistical differences were observed between the GCF and BGC module ordination analyses and, thus, we only present the results from the GCF analysis. Finally, the Jaccard dissimilarity matrix was used as input for a permutational multivariate analysis of variance (PERMANOVA) for 999 permutations.

To compare *Salinispora* biosynthetic potential to other genera, we searched the PATRIC v3.5.39 database (70) for a range of genera spanning the bacterial domain. Representative or reference genomes (as designated in the PATRIC database) were downloaded for each genus (N=582 genomes). The phylum Actinobacteria and family Micromonosporaceae were extensively sampled for direct comparisons to *Salinispora*. For each downloaded genome, we assigned protein-coding regions with Prokka, predicted BGCs with antiSMASH v5.0, and calculated the percent of the genome dedicated to specialized metabolism as the number of bp in all identified BGCs/genome length (bp).

**BGC evolutionary analyses**

For nine BGCs (Table S2), we extracted the BGC modules containing the respective MIBiG reference BGC from the network. Each BGC within the module was validated by screening for core biosynthetic genes known to be integral for product biosynthesis (Supplementary Table S2). In addition, we performed a second verification for each BGC by applying a targeted search with BiG-SCAPE using the MIBiG reference BGC as the query BGC (distance=0.7). The BiG-SCAPE output was used for BGC visualizations.

For phylogenetic analyses, we aligned all verified BGCs with Mugsy (71) to generate a nucleotide alignment as the input for phylogenetic analysis using RAxML under the GTRGAMMA model with 100 replicates. The phylogenetic distances for each BGC were compared to divergence times calculated from the molecular clock analyses for the same strain pairs. The BGC phylogeny was used as input for the event-inference parsimony model in Notung (72) by first rooting the phylogeny to minimize duplication and loss events for the top 10 parsimonious situations and reconciling each situation using a duplication (D=2), transfer (T=3), and loss (L=1) score to achieve the most parsimonious outcome. Both the phylogeny and nucleotide alignment were then used to calculate the relative influence of recombination and mutation with ClonalFrameML (73). The r/m values (relative contribution of recombination to mutation) were calculated by incorporating the length and genetic distance of the recombining fragments. As a comparison to the genome, we also aligned 28 representative genomes across the nine species and computed recombination metrics under identical parameters. To construct whole-genome recombination networks, we identified recent transfer events across all 118 *Salinispora* genomes using PopCOGenT (74), which uses a null model of sequence divergence to calculate a length bias to estimate recombining genomic segments.

We estimated the synonymous to nonsynonymous substitution rates (dN/dS) of shared biosynthetic genes in each of the nine target BGCs. Specifically, we clustered all genes in each BGC using CD-HIT-EST (75) at an 80% sequence identity threshold with the most similar clustering algorithm. Conserved BGC genes (i.e., genes present in every genome encoding the BGC) were aligned using Clustal Omega and used to calculate dN/dS ratios and nucleotide diversity using the PopGenome package (76) in R. As a comparison to the whole genome, we calculated nucleotide diversity of the 2011 core genes used to construct the core genome phylogeny above.

Salinosporamide AT domains were extracted from all strains containing the *sal* BGC (N=30). Specifically, we extracted all protein domains within the BGC and filtered based on the “PKS_AT” annotation from the *salA* gene. All AT domains were compared against the reference domain in *S. tropica* strain CNB440 (MIBIG: BGC0000145) with BLAT (77) for further confirmation. Both AT_1_ (extender) and AT_L_ (loading) were initially included by aligning all AT domains with Clustal Omega and constructing a phylogeny with RAxML under the PROTGAMMABLOSUM62 protein model. After confirming AT_1_ and AT_L_ domains were phylogenetically distinct, we reran the phylogenetic analyses only for the AT_1_ domain under identical parameters. For reference, AT domains from the *Streptomyces* BGCs that encode cinnabaramide (MIBIG: BGC0000971) and avermectin were also included, the latter had the closest protein homology as identified by SWISS-MODEL (78). Finally, we generated a protein structure homology-model with the reference AT_1_ domain in *S. tropica* against the avermectin (PDB:4RL1) template to calculate the most likely secondary structure under the DSSP program. All molecular graphic representations were visualized with PYMOL (https://pymol.org/2/).

**Strain cultivation and extraction**

We grew 30 representative *Salinispora* strains in A1 media with the following formulation: 10 g/L soluble starch, 4 g/L yeast extract, 2 g/L peptone, 10 g/L CaCO_3_, 22 g/L Instant Ocean dissolved in DI water. Cultures were supplemented with 5 mL/L KBr (20 g/L), 5 mL/L Fe_2_(SO_4_)_3_ (8 g/L) in 50 mL of A1 medium with metal springs added to reduce clumping and shaken at 205 RPM at 28°C. After a week of preculture, we assigned strains as fast- or slow-growing based on visual cell densities. For the slow-growing strains, we transferred 0.5 mL of culture to new media flasks (identical formulation except CaCO_3_ was omitted) and allowed another week of growth before the final transfer. All strains were subsequently transferred in triplicate to A1 media (1% v/v inoculum) supplemented as described above and shaken at 215 RPM at 30°C. On the 4th day of cultivation, 1 g of activated Amberlite XAD-7 resin was added to each flask (30 strains × 3 replicates=90 culture flasks + 3 media controls) to allow for passive adsorption of extracellular molecules. Based on visual evidence of sporulation and/or cell densities, cultures were extracted on the 7th or 10th day of cultivation using a 1:1 ethyl acetate solvent mixture. Specifically, 50 mL of ethyl acetate (EtOAc) was added directly to the culture flasks and allowed to shake for 2 h at 230 RPM. After shaking, the aqueous layer was discarded, and the organic layer collected. We removed any remaining water in the organic layer with Na_2_SO_4_ and, subsequently, filtered to remove resulting solids (Fisher Scientific; 0979014G). Finally, extracts were dried by rotary evaporation in vacuo, resuspended in HPLC-grade EtOAc (Fisher Scientific; E195-4), and dried under nitrogen gas for storage at -20°C until later processing.

**Metabolomics and mass spectrometry**

Crude extracts were resuspended in HPLC-grade methanol to a concentration of 1 mg/mL and filtered by centrifugation (0.2 µm). Samples were analyzed via liquid chromatography-tandem mass spectrometry (LC-MS/MS) with 1 μL injections into an Agilent 1290 High Performance Liquid Chromatography (HPLC) system coupled to an Agilent Accurate Mass QToF spectrometer. For targeted analysis of the salinosporamides, 5 μL injections were also analyzed to ensure MS/MS fragmentation of the targeted compounds. Standards for lomaiviticin C, lymphostin and neolymphostin A, rifamycin W, salinipostins C, E, and H, salinosporamides A and B, and staurosporine were also analyzed. Parameters were set to allow a flow rate of 0.75 ml/min through a Phenomenex Kinetex C18 reversed-phase column (5 μm, 100 x 4.5 mm; Phenomenex, Torrence, CA, USA) for chromatographic separation. For reverse phase chromatography, mobile phase of solvents A (water with 0.1% formic acid (FA) [v/v] and B (acetonitrile with 0.1% FA [v/v]) were used with conditions as follows: 0-4 min (95% A, LC Stream diverted to waste), 4-24 min (95% to 0% A), 24-26 min (0% A), 26-26.5 min (0% to 95% A), 26.5-30 min (95% A). Positive ion mode data was collected in profile mode. MS1 data was acquired from 100 – 1,700 m/z with the top 5 most abundant precursor ions selected for MS2 fragmentation using a fixed collision energy of 30 eV and acquired from 50 – 1,700 m/z. Two MS2 scans were acquired per second and ions were excluded from fragmentation for 30 seconds if acquired 3 times. MS data was collected with source parameters: nebulizer gas (nitrogen) at 35 psig, drying gas flow of 11 l/min, capillary voltage of 3000 V, gas temperature of 300° C, acquiring 3 spectra per second. All datafiles were converted to mzXML using the MSConvert tool in the Trans-Proteomic pipeline (79) for further downstream analysis. Paired genomic and metabolomic datasets for each strain are available on the Paired Omics Data Platform (80).

Datafiles were preprocessed with MZmine v2.37.1 (81) with ion identity networking. Preprocessing parameters were set based on raw MS data, including the centroided masses at a noise level of 1E3 for MS1 and 3E1 for MS2 scans. The features table was produced by comparing chromatograms with the ADAP chromatogram builder, deconvoluted using the local minimum search, and isotopic peaks removed. Individual deconvoluted chromatograms were joined into an aggregated peak list with gap filling using an intensity tolerance of 10%, m/z tolerance of 0.0 *m/z* or 10 ppm and a retention time tolerance of 0.2 minutes. For the ion identity network, the MetaCorrelate function was used with a minimum of five data points: two minimum data points on edge, a minimum feature shape correlation of 85%, and an *m/z* tolerance of 0.0 *m/z* or 10 ppm. All features were manually checked with a minimum height of 1E3 for the following ions: [M+H] ^+^ (1.0073), [M+Na]^+^ (22.9892), [M+K]^+^ (38.9632), [M+2H] ^2+^ (2.0146), [M+Ca]^2+^ (39.9615), [M+Fe] ^2+^ (55.9338), [M+Fe-H]^+^ (54.9266), [M+Fe]^3+^ (55.9333), and [M+Fe-2H]^+^ (53.9187). For untargeted metabolomic analysis, mass spectrometry data files from the 1 μl injections were used to generate an MS1 feature table using MZmine adapted for Ion Identity Networking. Data preprocessing parameters were generated from careful analysis of raw MS data. A final peak list was produced using the ADAP Chromatogram builder, quality filtering, chromatogram deconvolution, isotopic peak grouping, join alignment, gap filling, further quality filtering of gap filled peaks, and ion identity networking.

We compared metabolic profiles of 30 *Salinispora* strains using the MS1 molecular feature table. First, we normalized the 3575 identified features across all samples (90 strains, 3 media controls, 1 column wash, and 11 standard compounds) using a log_2_ transformation with pareto scaling. A preliminary Principal Components Analysis ordination plot was generated for the 105 samples to assess data quality compared to standards and control samples. After manually removing all features found in standards and wash samples and subtracting feature intensities found in media controls, the remaining 3168 features were normalized as above. Next, we averaged the intensities across the triplicate samples for each strain, generated a Euclidean distance matrix, and performed a permutational multivariate analysis of variance (PERMANOVA (82)) with species as a fixed effect for 999 permutations under a reduced model. Finally, the average strain feature data were used to generate a final Principal Components Analysis ordination plot.

The MS2 classical molecular network was initially used to identify the nine compounds of interest and their analogs by extracting clusters containing either GNPS library matches or network nodes containing the standard compounds (Table S3). For sporolide, salinichelins, and salinilactams, where no standards were available, we manually curated the extracted ion chromatogram (EICs) at a maximum of 10 ppm error for the theoretical *m/z* of the molecules [M+H]^+^ and their analogs, and verified production if both the [M+H]^+^ and [M+Na]^+^ adducts were present in the MS1 chromatograms (Figure S14). When possible, these masses and retention times were linked back to the MS2 network. Based on the results from the MS2 network and EICs, theoretical *m/z* [M+H]^+^ values for all analogs were correlated with the MS1 feature table based on *m/z* (∆ppm=2.2) and retention times (Table S3).

MS analysis of the nine compounds corresponded to 28 features in the molecular feature table. After removing features found only in standards (i.e., salinipostins), the remaining 25 features were extracted and normalized using a log transformation. The 25 feature intensities were averaged across triplicate samples per strain and validated using MS2 spectra from the classical molecular network (see above). Specifically, analog production was only confirmed if strains had intensity values corresponding to the MS1 feature table and were also identified in the MS2 classical molecular network. Using the confirmed analog production intensities, a PERMANOVA was performed using a Euclidean distance matrix with species as a fixed effect for 999 permutations under a reduced model. Finally, we performed a hierarchical clustering analysis on the Euclidean distance matrix for a heatmap visualization of analog production and added the presence or absence of each of the BGCs corresponding to the nine molecules. All statistical analyses were performed in R.

For targeted metabolomics of the salinosporamides, the 5 μL injections (at 1 mg/mL) were processed as above to generate a classical MS2 molecular network. We extracted the salinosporamide cluster using both the GNPS library matches and the standards for salinosporamides A and B (Supplementary Figure S11). To calculate the relative production of each salinosporamide analog, we manually curated the extracted ion chromatogram (EICs) at a maximum of 10 ppm error for the theoretical *m/z* of the molecules [M+H]^+^ and their adducts. We extracted intensity values from the chromatograms if the [M+H]^+^, [M+Na]^+^, and in the case of salinosporamide A, the presence of the Cl isotope signature (at *m/z*=316.114) were all above the 1E3 noise threshold level. Production was confirmed if the samples were also identified in the salinosporamide MS2 network.
